# ABaCo: Addressing Heterogeneity Challenges in Metagenomic Data Integration with Adversarial Generative Models

**DOI:** 10.1101/2025.09.22.677692

**Authors:** Edir Vidal, Angel L. Phanthanourak, Atieh Gharib, Henry Webel, Juliana Assis, Sebastián Ayala-Ruano, André F. Cunha, Alberto Santos

## Abstract

The rapid advancement of high-throughput metagenomics has produced extensive and heterogeneous datasets with significant implications for environmental and human health. Integrating these datasets is crucial for understanding the functional roles of microbiomes and the interactions within microbial communities. However, this integration remains challenging due to technical heterogeneity and the inherent complexity of these biological systems. To address these challenges, we introduce ABaCo, a generative model that combines a Variational Autoencoder (VAE) with an adversarial discriminator specifically designed to handle the unique characteristics of metagenomic data. Our results demonstrate that ABaCo effectively integrates metagenomic data from multiple studies, corrects technical heterogeneity, outperforms existing methods, and preserves taxonomic-level biological signals. We have developed ABaCo as an open-source, fully documented Python library to facilitate, support and enhance metagenomics research in the scientific community.

## 1 Introduction

Advances in sequencing technologies have enabled population-scale metagenomic studies in diverse environments. Several public platforms host extensive collections of microbiome data, enabling a wide range of downstream analyses. For instance, the MGnify platform offers more than 450,000 annotated genomes in Metagenome-Assembled Genomes (MAGs) catalogues, and also provides automated pipelines for the assembly, analysis, and archiving of metagenomic and amplicon sequencing data [1]. Similarly, DOE-JGI’s IMG/M offers tools for curation and comparative analysis of microbial genomes and metagenomes [2]. Large collaborative efforts, such as the Human Microbiome Project (HMP), have characterized the healthy human microbiome by analyzing samples from hundreds of individuals [3]. More recently, The Open MetaGenomic dataset (OMG) aggregated over 3.1 terabases (≈3.3 billion coding sequences) from IMG/M and MGnify [4]. These initiatives demonstrate the scale and diversity of modern microbiome datasets and highlight the potential for atlas-level integrative analyses.

Despite these advances, integrating data from different projects and platforms remains challenging due to technical heterogeneity. Figure 1a illustrates the main sources of variation encountered in the integration of metagenomic datasets. In particular, differences in sequencing technologies (e.g., 16S rRNA amplicon vs shotgun sequencing, or Illumina vs long-read platforms) introduce cross-platform heterogeneity that complicates integration. Additionally, variations in laboratory protocols (DNA extraction, library preparation, etc.) can lead to systematic biases [5]. Even minor protocol variations in metagenomics can substantially alter observed community profiles. These unwanted sources of variation, known as *batch effects*, are unrelated to the biological factors of interest [5]. Batch effects can distort true biological signals, thereby undermining conclusions drawn from cross-study analyses. Correcting for such technical heterogeneity is critical in large-scale metagenomic studies.

**Figure 1:**
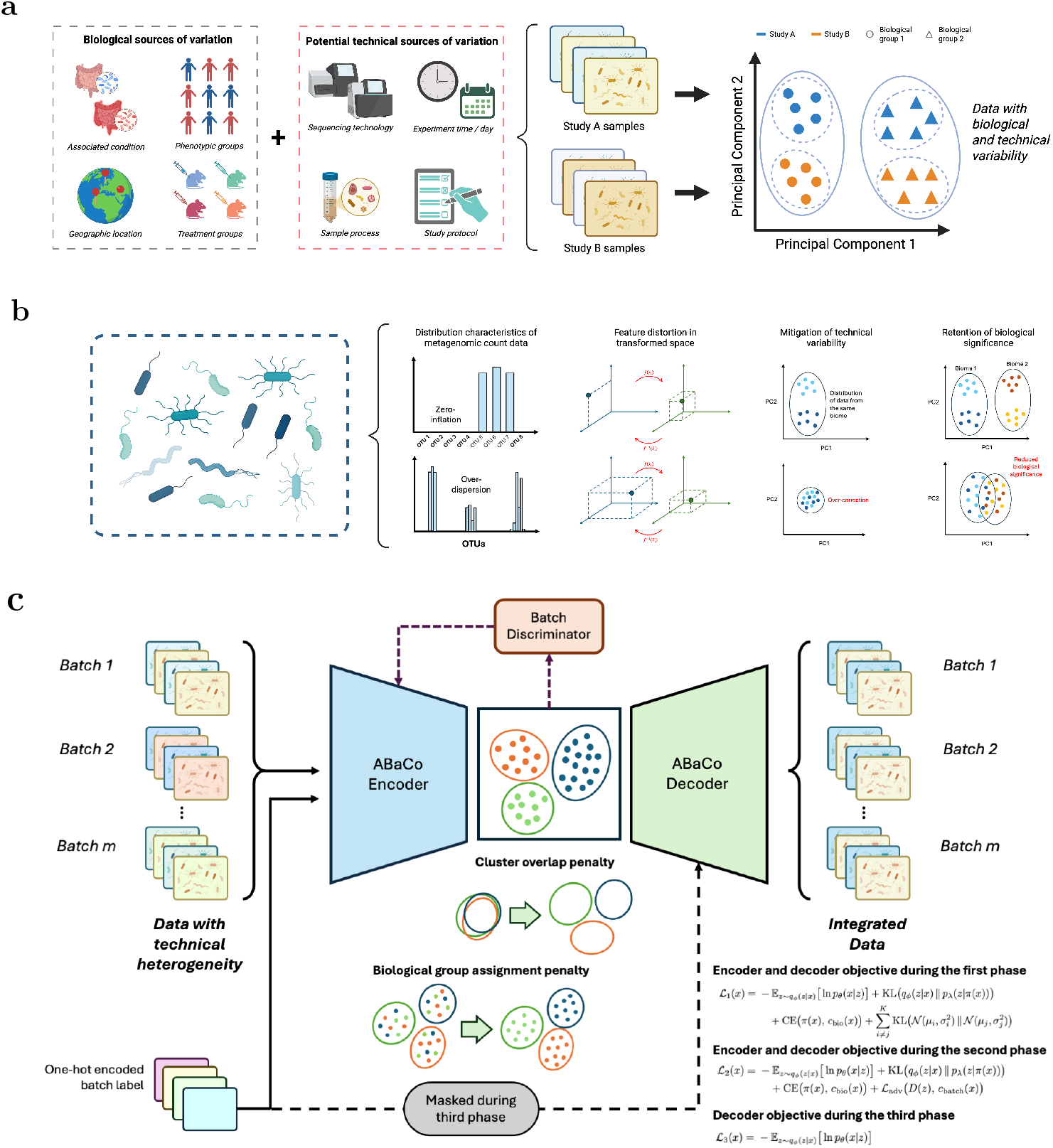
Schematic summary of variation sources, integration challenges and the ABaCo architecture. **(a)** Biological and technical main sources of variation and their effect on sample clustering in metagenomic studies. **(b)** Integration challenges: count distributions (zero-inflation, over-dispersion), transformation distortions, and the trade-off between technical mitigation and biological retention. **(c)** ABaCo architecture, illustrating the VAE, batch discriminator, and the cluster overlap and biological group assignment penalties within a phased training scheme; loss terms are noted alongside the model. Figure created in BioRender.

Current batch correction methods face limitations addressing metagenomic technical heterogeneity (Figure 1b). Classical batch effect correction methods, originally developed for gene expression data, can be applied to microbiome abundance tables. Parametric linear model approaches such as ComBat [6] and limma [7] employ empirical Bayesian frameworks to adjust for batch effects across features. These methods effectively remove global shifts under the assumption of normality and are widely used in transcriptomics. However, metagenomic data rarely meet these assumptions: read counts are highly sparse and compositional, representing relative abundances that sum to a constant [8]. Ignoring this data structure can lead to misleading inferences [8].

More recent tools designed specifically for metagenomic applications aim to capture the complexity of these datasets better. For example, ConQuR [9] employs a two-part conditional quantile regression model to account for zero-inflation and over-dispersion in microbial read counts, yielding batch-corrected values that preserve the complex count distribution. Another approach, PLSDA-batch [10], uses a multivariate non-parametric method based on partial least squares discriminant analysis. It estimates latent components associated with both biological and technical variation, then subtracts the batch-related components to remove biases while preserving biological differences. However, low-frequency taxa and nonlinear confounding factors can impair these methods’ accuracy and complicate downstream interpretation [11].

Large-scale microbiome studies such as metagenomics and metabarcoding demand robust batch correction, yet traditional methods may not fully address the sparse, compositional nature of microbiome counts. While specialized tools incorporate data-specific modeling, the field is increasingly moving towards deep learning solutions that can leverage complex dataset structure with fewer assumptions. Examples from single-cell work include ABC [12] and scDREAMER [13], which use adversarial alignment and showcases how training an autoencoder with auxiliary cell-type classifiers along with adversarial alignment can effectively remove batch effects while preserving biological labels. Most recently, the VampPrior Mixture Model (VMM) [14] has been employed as a prior distribution to the widely used scVI model [15], enhancing its batch correction capability while better retaining biological information in single-cell datasets. These approaches highlight how deep generative modeling can flexibly extract the complexity of data distributions and disentangle technical variation from biological signal.

The methodological parallels in data integration on single-cell transcriptomics and metagenomics are driven by shared challenges: high dimensionality, extreme sparsity of features, and complex confounded technical factors. Here, we introduce **ABaCo**, a generative adversarial framework designed for metagenomic batch correction. ABaCo integrates recent advances from single-cell transcriptomics, such as adversarial training and clustering-based priors and adapts them to the challenges of microbial count data horizontal integration. We demonstrate that ABaCo not only achieves state-of-the-art performance but also removes technical variation between studies while preserving biological signals, improving downstream analyses of microbiome profiles.

## 2 Results

### 2.1 Datasets overview

We benchmarked ABaCo (Figure 1c) against state-of-the-art methods across scenarios to evaluate performance and robustness. We created simulated count datasets to establish a controlled environment with known ground-truth biological groups. Additionally, we assessed ABaCo’s performance on horizontal data integration tasks using a series of independent datasets from different sources (case studies), including anaerobic digestion assays, Inflammatory Bowel Disease (IBD) patient samples, and sewage metagenomic profiles. For reproducibility and for the figures shown, we generated a representative simulated replicate using a predefined seed (42). All benchmarking and statistical results, however, are computed over 50 independent simulated replicates per scenario. As shown in Figure 2a, both the simulated datasets and the case studies exhibit a highly skewed proportion of zero counts, indicating widespread zero inflation. This is an inherent characteristic of metagenomic data, where a large subset of taxonomic features are absent in most samples, resulting in a long left tail in the distribution of non-zero proportions.

**Figure 2:**
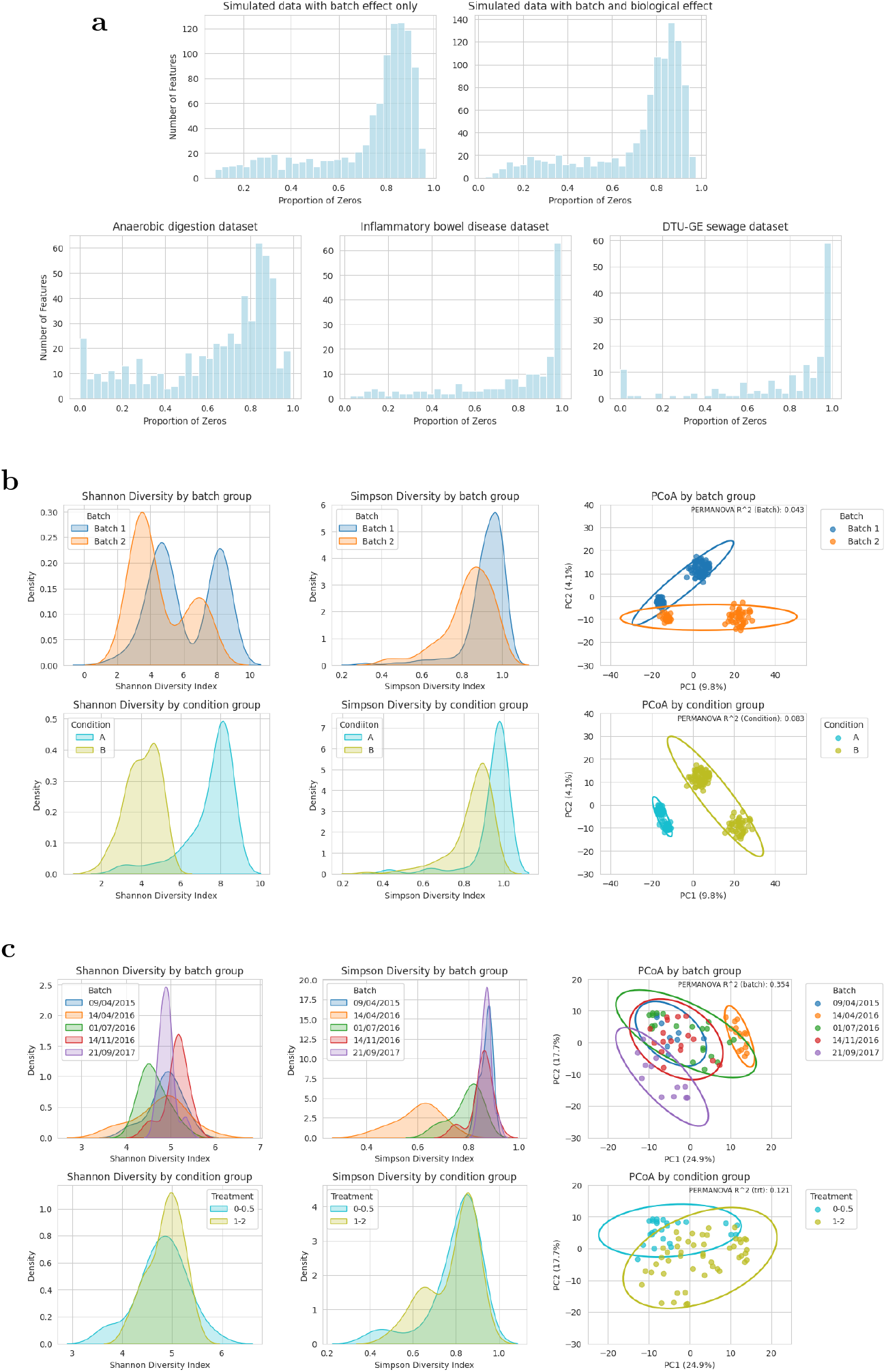
Comprehensive comparison of data sparsity, alpha diversity and beta diversity across simulated and case study datasets. **(a)** Histograms of the proportion of zero counts per taxonomic feature for simulated and case study datasets. **(b–c)** Kernel density estimates of Shannon and Simpson alpha diversity (left, center) and PCoA on Aitchison distances (right), shown by batch (top) and biological group (bottom) for the simulated dataset (b) and the anaerobic digestion case study (c).

#### 2.1.1 Simulated datasets

We generated two complementary sets of simulated datasets sampling from the zero-inflated negative binomial (ZINB) distribution to evaluate model performance. The first scenario included only batch effects with no biological differences between groups, while the second scenario contained both biological signal and batch differences. Each replicate consisted of 200 samples and 1,000 features, divided into two biological groups (50:50) and two batches (60:40). To focus evaluations on sampling variability, the parameters of the sampling distribution were kept constant across replicates. We created 50 independent replicates for each scenario and used the resulting counts to evaluate the model’s batch correction performance and consistency.

Figure 2b illustrates the distributions of Shannon and Simpson alpha diversity indices stratified by batch and biological groups, along with the beta diversity Aithchison distances Principal Coordinate Analysis (PCoA) for the simulated data with batch and biological effects. Both alpha diversity indices display a clear shift concordant with the imposed batch and biological effects: the two biological groups differ in median alpha diversity and in the spread of values, as do the two batch groups. Additionally, the PCoA reveals strong partitioning on both batch and biological effects, with the biological effectbeing the primary driver, as indicated by the PERMANOVA R^2^. These results indicate that the simulated biological signal is the primary source of compositional differences, while the simulated batch effect contributes a smaller, yet visually evident, source of variation. The same illustration can be found on Supplementary Figure 1 for the simulated dataset with batch effect only.

#### 2.1.2 Case studies datasets

To evaluate ABaCo’s performance across diverse case studies, we selected datasets from MGnify and a benchmark dataset designed for assessing batch effects in microbiome data. An overview of these datasets is provided in Table 1, detailing the batch and biological groups. Notably, the anaerobic digestion and DTU-GE sewage datasets are semi-balanced across batches, with nearly equal samples sizes per batch, whereas the IBD dataset has a high disparity among batch groups. Both the DTU-GE sewage and IBD datasets show high variability in the relative abundance of the most dominant taxa. Importantly, the separation between biological groups is not consistent across batches, indicating that batch-specific effects may influence the biological inference. To assess the consistency of ABaCo, we trained the model 50 times on each dataset with different initialization, using a defined seed (42) only for the first training instance.

**Table 1:** Overview of datasets, batches, biological groups, and sample counts for every case study. For each dataset the table reports the number of taxa, the batch group identifier (date, accession or pipeline), the biological-group labels present in that batch (treatment, phenotype or location) and the per-group sample counts; the right-most column gives the total number of samples in each batch and dataset.

| Dataset | Taxa | Batch | Biological group | Samples | Batch total |
| --- | --- | --- | --- | --- | --- |
| Anaerobic digestion | 567 | 09/04/2015 | 0–0.5 | 4 | 9 |
|  |  |  | 1.0–2.0 | 5 |  |
|  |  | 14/04/2016 | 0–0.5 | 4 | 16 |
|  |  |  | 1.0–2.0 | 12 |  |
|  |  | 01/07/2016 | 0–0.5 | 8 | 21 |
|  |  |  | 1.0–2.0 | 13 |  |
|  |  | 14/11/2016 | 0–0.5 | 8 | 17 |
|  |  |  | 1.0–2.0 | 9 |  |
|  |  | 21/09/2017 | 0–0.5 | 2 | 12 |
|  |  |  | 1.0–2.0 | 10 |  |
| Inflammatory Bowel Disease | 435 | PRJNA389280 | Crohn’s Disease (CD) | 180 | 348 |
|  |  |  | Ulcerative Colitis (UC) | 99 |  |
|  |  |  | non IBD | 69 |  |
|  |  | PRJNA398089 | Crohn’s Disease (CD) | 85 | 176 |
|  |  |  | Ulcerative Colitis (UC) | 46 |  |
|  |  |  | non IBD | 45 |  |
| DTU-GE sewage | 162 | pipeline 4.1 | Canada | 11 | 56 |
|  |  |  | USA | 30 |  |
|  |  |  | Zambia | 7 |  |
|  |  |  | Australia | 8 |  |
|  |  | pipeline 3.0 | Canada | 13 | 73 |
|  |  |  | USA | 40 |  |
|  |  |  | Zambia | 10 |  |
|  |  |  | Australia | 10 |  |

**Anaerobic digestion study:** this benchmark dataset, used for microbiome batch effect assessment [5], comprises anaerobic digestion of organic matter under varying phenol concentrations [16]. The dataset includes 567 Operational Taxonomic Units (OTUs) identified from the microbiota of 75 samples collected in batches on different dates, which is the main source of technical variation. Figure 2c provides an overview of the alpha and beta diversity of the dataset. The distribution of Alpha diversity indices indicates that batch effects are the primary source of variation, as evidenced by lower overlap between batches compared to biological groupings. The PCoA further confirms this effect, showing that the biological effect remains significantly different between the two treatment groups despite the batch variation.

**Inflammatory bowel disease projects:** This case study comprises two different studies [17] [18] from Harvard T.H. Chan School of Public Health, available in MGnify, which analyze the microbiome dynamics of 524 patients with inflammatory bowel disease (IBD) and non-IBD controls. Within the IBD and non-IBD patient groups a total of 435 genus-level taxa were identified, with cross-study batch effect accounting for the main source of variation in the dataset (see Supplementary Figure 2). A distinction was made between Crohn’s disease (CD) and Ulcerative colitis (UC) for a more enriched analysis of the phenotypes taxonomic differences.

**DTU-GE Sewage global surveillance:** This study involves global sewage metagenome surveillance for pathogen and antimicrobial-resistance monitoring [19]. It includes point-prevalence metagenomic profiles from raw sewage collected at main sewer inlets of major cities worldwide. The data was processed using two different MGnify pipelines at the phylum level, introducing technical heterogeneity to the integrated dataset. For our analysis, we focused on samples from Australia, USA, Canada, and Zambia, as each of these countries contributed at least 10 samples. After filtering, 162 taxa were retained with each pipeline accounting for the main source of variation in the data (see Supplementary Figure 3).

### 2.2 Performance on datasets

For a thorough evaluation of our model, it is essential to acknowledge the challenges posed by the complexity and heterogeneity of batch effects in biological datasets [20]. To address these issues effectively, we adopted a holistic approach that evaluates both the efficacy of batch effect correction and the preservation of biological signals, reporting graphical representations and quantitative metrics.

To evaluate the robustness of our methodology, ABaCo was trained multiple times with different initializations for the case studies and with a defined seed (42) for the simulated datasets. The training hyperparameters and weights used for each scenario are detailed on the Supplementary Table 1-2. ABaCo was tested using two output distributions: zero-inflated negative binomial (ZINB) and negative binomial (NB). The hyperpameters for these two configurations were the same.

We also applied state-of-the-art methods to correct the datasets using normalized counts (BMC, ComBat, limma and PLSDA-batch) and raw counts (ComBat-seq and ConQuR) to compare the performance of ABaCo. Figure 3 summarizes the batch correction performance using various metrics for all the mentioned methods: kBET [21] (local mixing of batches), iLISI [22] (integration at the sample level), batch ASW [23] (cluster overlap for batch groups) and batch ARI [24] (agreement between batch labels and clusters). All these metrics are normalized, were values approaching 1.0 reflecting improved batch correction performance.

**Figure 3:**
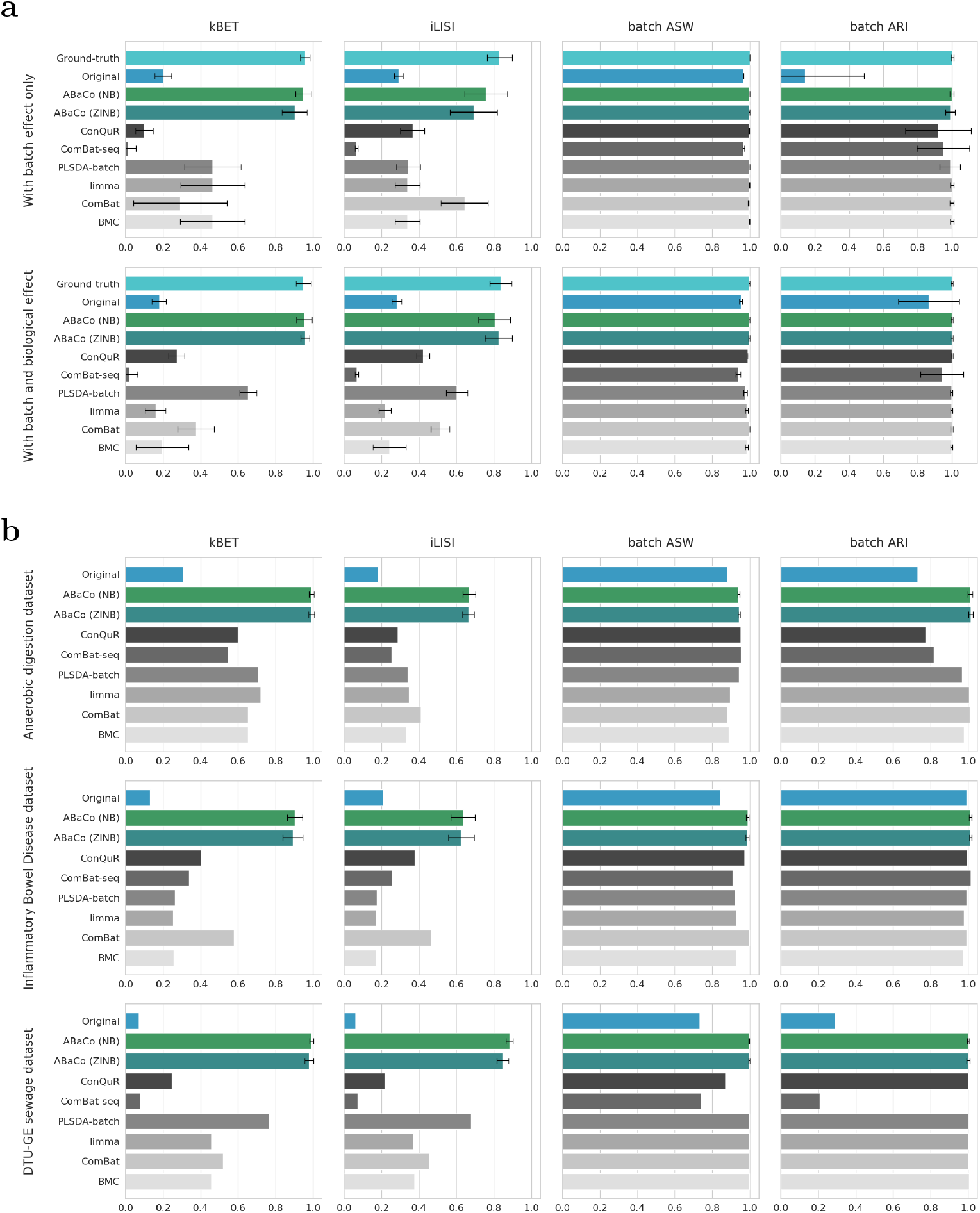
Performance of ABaCo and state-of-the-art methods on simulated and case study datasets. **(a)** In simulated datasets were the ground-truth is reported, ABaCo performs well both with and without biological effects, outperforming alternatives in local batch correction (kBET, iLISI) while maintaining global structure comparably (batch ASW and ARI). **(b)** In case studies, ABaCo likewise surpasses state-of-the-art methods (kBET, iLISI) and preserves global structure to a similar extent (batch ASW and ARI).

Figure 3a demonstrates that ABaCo, with both negative binomial (NB) and zero-inflated negative binomial (ZINB) distributions, achieves the best performance in batch effect correction compared to other methods for the kBET and iLISI metrics. For the batch ASW and batch ARI metrics, ABaCo performs similarly to the top methods. Notably, ABaCo with the ZINB distribution showed lower variance and higher mean performance in the kBET and iLISI metrics for the simulated data with both batch and biological effects.

In Figure 4a the PCoA showcases that ABaCo effectively integrates the batch groups of the simulated data containing both batch and biological effects. With the ZINB model, ABaCo achieves the closest alignment to the ground-truth PCoA plot, successfully mixing the batches and preserving the biological structure. ABaCo also handles batch effects without adding biological bias for the simulated data with batch effect only (see Supplementary Figure 4).

**Figure 4:**
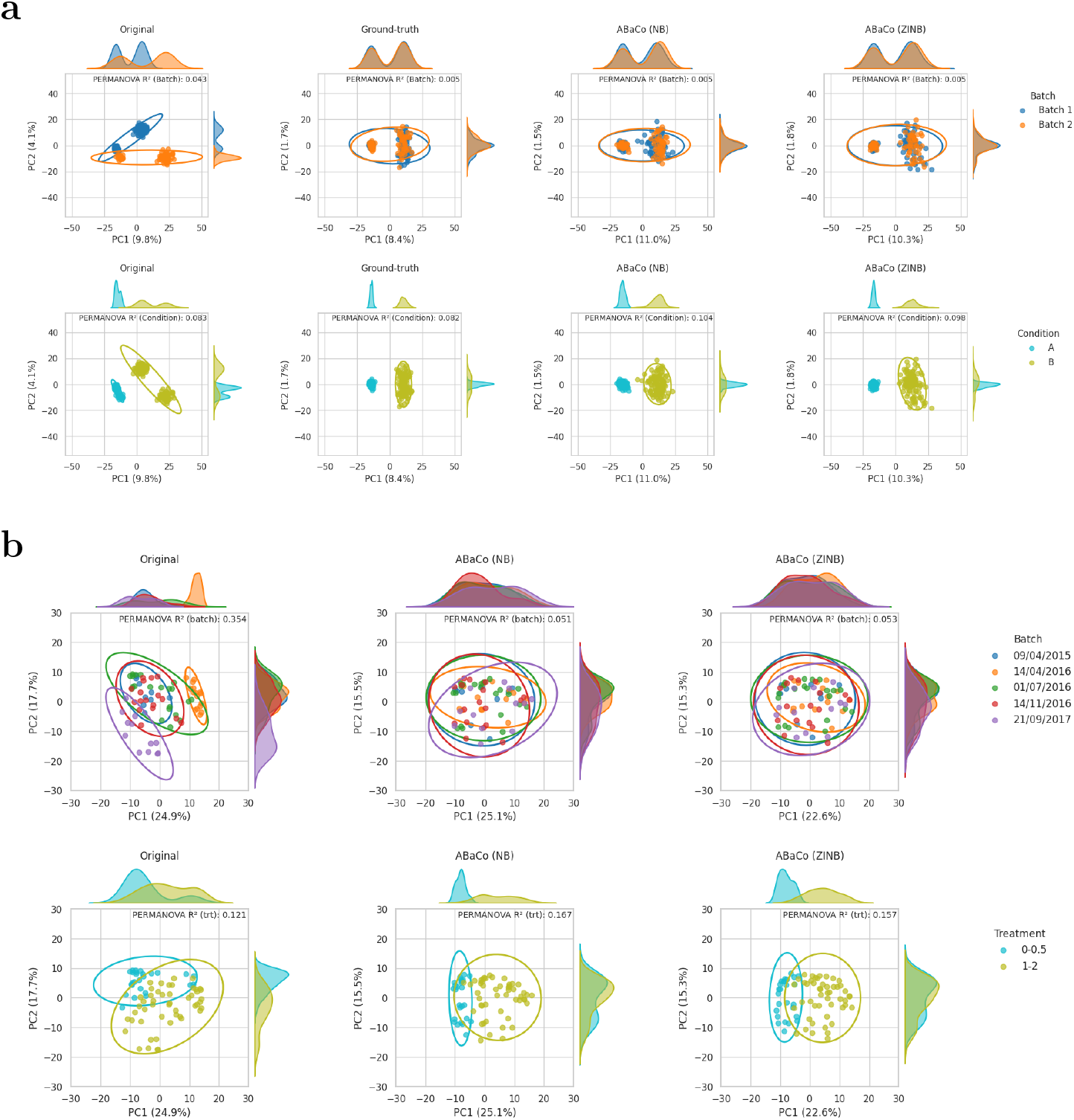
Principal Coordinates Analysis (PCoA) using Aitchison distance on datasets corrected with ABaCo (ZINB and NB models). **(a)** Simulated data containing both batch and biological effects; the ground-truth PCoA is shown for reference. **(b)** Anaerobic digestion case study. For each dataset, ABaCo mix batch groups while preserving biologically relevant group structure.

Regarding the case studies datasets, Figure 3b illustrates ABaCo’s performance compare to the other methods. ABaCo, with both NB and ZINB distributions, outperforms other methods in the kBET and iLISI metrics while preserving the biological signal, as indicated by comparable batch ASW and ARI metrics.

The benchmarking metrics demonstrate that ABaCo effectively addresses batch effects from multiple groups. This is also visible confirmed in the figures for the anaerobic digestion dataset (Figure 4b) and the other two case studies (Supplementary Figure 5-6), where PCoA plots show the batch effects corrected and the biological differences maintained. We did the same analysis using the state-of-the-art methods mentioned for all datasets (Supplementary Figure 7-11).

### 2.3 Robustness analysis

We assessed ABaCo’s robustness by tracking the most abundant taxa in each case study after batch correction across all training runs. Figure 5 reports the mean relative abundance of the five most abundant taxa, stratified by biological group. Statistical comparisons between groups were performed using the Kruskal–Wallis test (Supplementary Figure 12).

**Figure 5:**
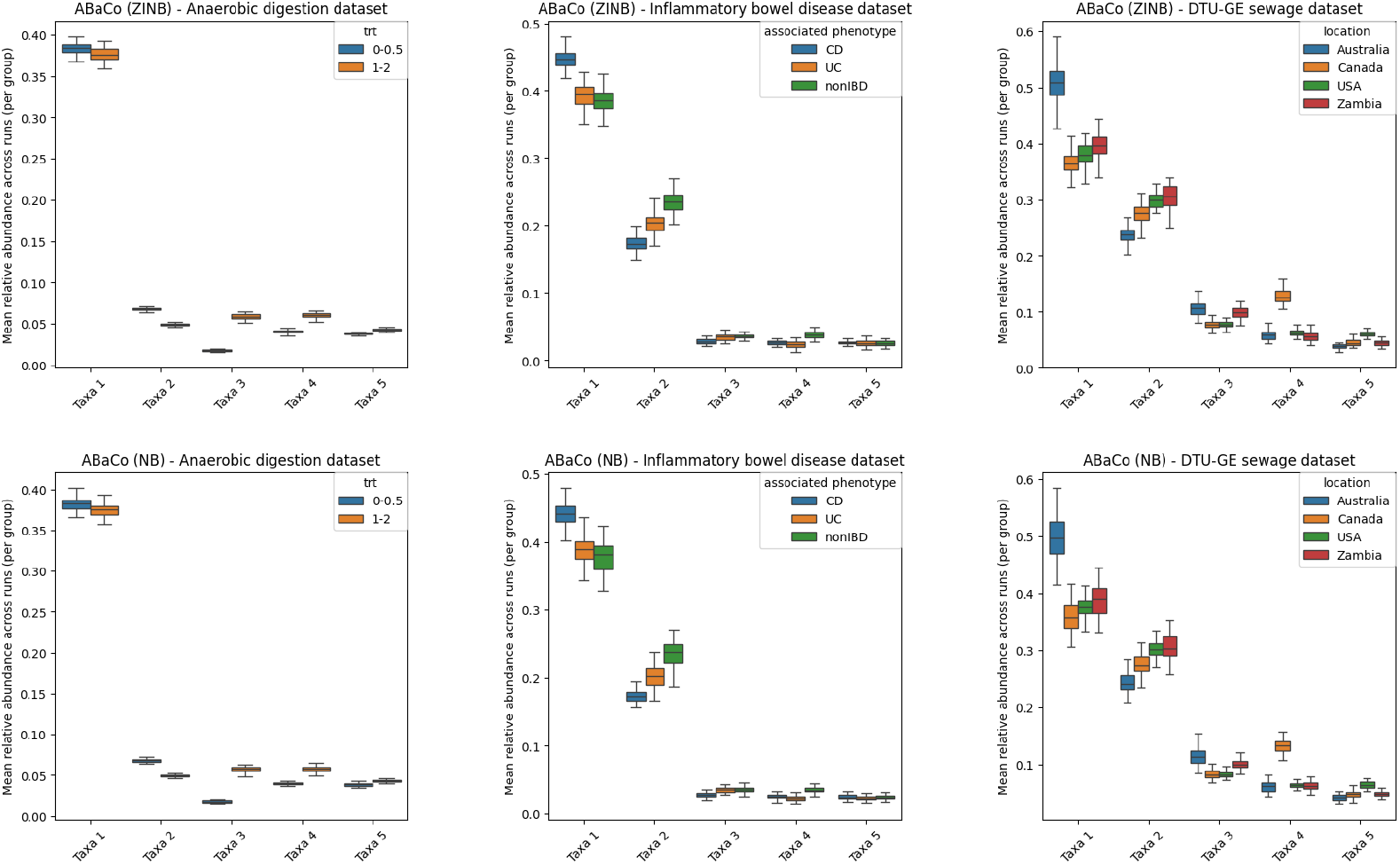
Box-plots of the five most abundant taxa mean (relative abundance) for all training iterations of ABaCo. Mean value is registered after every training iteration with the grouping based on condition (treatment group, associated phenotype, or geographic location).

For the anaerobic digestion case study, both ZINB and NB models return very low variance across runs for the top five taxa (variance range: 1.6 × 10^−6^ to 7.1 × 10^−5^), indicating consistent reconstruction of dominant taxa. In the IBD and DTU-GE datasets the most abundant taxa were likewise preserved: the two most frequent taxa together account for more than 50% of total counts in their respective datasets. Variance for the two most abundant taxa was higher than for lower-ranked taxa (IBD, taxa 1–2: 9.1 × 10^−5^ to 5.9 × 10^−4^; taxa 3–5: 1.2 × 10^−5^ to 3.3 × 10^−5^; DTU-GE, taxa 1–2: 2.0 × 10^−4^ to 1.7 × 10^−3^; taxa 3–5: 2.5 × 10^−5^ to 3.2 × 10^−4^). Overall, these results show that ABaCo reliably preserves the dominant taxonomic features across runs, with larger variance concentrated in the most abundant taxa.

## 3 Discussion

We have presented ABaCo, a generative adversarial framework that combines a variational autoencoder with adversarial discrimination and count-based distributions (NB) that account for zero-inflation (ZINB) to mitigate technical heterogeneity while preserving biological signals in horizontal metagenomic integration. Across both simulated and real-world case studies, ABaCo consistently reduced batch-associated structure, as evidenced by high-scoring kBET and iLISI metrics, and preserved biological separation, as demonstrated by biologically informed PCoA plots and test statistics. Notably, the ZINB variant yielded more stable performance (lower variance across runs) in the simulated scenario with both batch and biological effects, suggesting that explicitly modeling zero inflation improves robustness for sparse microbiome profiles.

Robustness analyses, which assess consistency across runs or variance in output behavior, are still relatively uncommon for generative models. However, they are crucial for evaluating model reliability beyond best-case scenarios, diagnosing sources of instability, and setting realistic expectations for deployment. To this end, we conducted such analyses and found that, on average, run-to-run reproducibility of the most abundant taxa after batch correction was high across datasets, indicating highly consistent reconstruction of dominant community members. These results support the claim that ABaCo reduces technical heterogeneity without erasing biologically meaningful variation. This pattern also suggests that larger absolute abundances amplify run-to-run variance even when overall taxonomic structure is preserved, and that highly heterogeneous cohorts (e.g. IBD) can produce a small number of extreme outlier runs. IBD samples are typically more variable across individuals and studies, and the combination of high biological heterogeneity across batches may amplify sensitivity to initializations or hyperparameters. Therefore, we recommend that analyses of similarly heterogeneous atlases include multiple training replicates, inspection of low-agreement runs, and reporting of both mean tendency and tail statistics rather than relying solely on averages.

Explicitly modeling count distributions proved beneficial for performance on sparse microbiome data, with the ZINB model demonstrating lower variance in challenging simulated settings, aligning well with observed zero-inflation patterns. Additionally, the adversarial discriminator effectively removed provenance signals in the latent space, highlighting its strengths in mitigating batch effects. The balance between adversarial and biological-preservation losses is crucial, as it influences the trade-off between batch removal and the conservation of biological signals. To optimize this trade-off, we recommend using a small validation set when applying ABaCo to new datasets. This approach allows for fine-tuning the weight of the adversarial loss, ensuring robust batch effect removal while preserving biologically meaningful variations.

ABaCo effectively models observed counts and zero inflation, providing a robust framework for analyzing sparse metagenomic data. While it does not intrinsically enforce compositional constraints, thoughtful preprocessing choices such as filtering rare taxa, library-size normalization, or compositional transforms can improve downstream analyses and should be consistently reported to ensure reproducibility. In addition, adversarial training can be sensitive to optimizer schedules and discriminator–encoder balance. By implementing carefully staged learning rates, we reduced mode collapse, demonstrating ABaCo’s adaptability to large datasets. Furthermore, the latent clustering prior helped preserve biological group structure in our experiments, indicating its potential for maintaining meaningful biological signals. Future work could focus on improving the interpretability of latent features, linking them to environmental covariates or phylogenetic structures to gain deeper biological insights.

To further enhance ABaCo’s capabilities, future developments could incorporate explicit compositional models to boost biological fidelity. Prior extensions could increase latent expressivity and better decouple batch effects from biological signals. For instance, addressing the inconsistency of the latent space which is common on VAEs could improve robustness in highly heterogeneous cohorts. Expanding the evaluation to include targeted downstream analyses presents a natural next step to demonstrate the practical benefits of ABaCo’s batch correction. Incorporating analyses such as differential abundance with FDR control, network inference, and functional profiling would help show how batch correction influences biological conclusions. We note that some of these evaluations are already possible with our outputs, but a full biological interpretation of the results is beyond the scope of the current paper. Nonetheless, we anticipate that future studies leveraging ABaCo should reveal enhanced biological relevance and actionable insights, paving the way for a more reliable and impactful microbiome research.

ABaCo provides a flexible, deep generative approach for the horizontal integration of metagenomic studies, striking a balance between batch removal with biological conservation. The framework demonstrates strong performance across multiple benchmarks and real datasets. When applied with appropriate preprocessing and systematic validation, can facilitate more reliable atlas-level microbiome analyses. To promote adoption, reproducibility, and accessibility, we developed ABaCo as an open-source Python library available via PyPI for easy installation and integration into workflows. Its comprehensive documentation enables rapid implementation of batch correction and allows other researchers to build on our work and contribute to further develop the model.

## 4 Methods

### 4.1 ABaCo framework

ABaCo is an adversarial generative framework trained in three sequential phases. In the first phase, a variational autoencoder (VAE) with a variational prior distribution is trained with batch labels supplied to both the encoder and the decoder. This setup also contains two penalization terms – a cross-entropy loss for biological group assignment, and a pairwise KL-divergence between the Gaussian components of the prior distribution – to structure the latent space according to biological groups, thereby promoting biologically meaningful clustering. In the second phase, the prior distribution parameters are set, and adversarial training is applied. The discriminator receives latent embeddings as input and backpropagates a gradient to the encoder. This adversarial setup encourages the encoder to remove batch-specific information from the latent representation while retaining meaningful biological signal. In the third phase, the decoder is trained with batch labels gradually masked. This ensures that the outputs are no longer dependent on batch-specific features. An overview of the model architecture can be found in Figure 1c.

#### 4.1.1 VAE setup

ABaCo uses the architecture of a regular VAE: an encoder that maps the data into a latent space, a prior distribution that regularizes the latent representation, and a decoder that reconstructs data points back into the original input space [25]. The encoder receives an input data point and outputs the parameters of a Mixture-of-Gaussian (MoG) distribution (means, variances, and mixing probabilities). A latent point is then obtained by first sampling a component from the Categorical distribution using the mixing probabilities, and then sampling from the corresponding Gaussian using the mean and variance. For the prior, ABaCo employs the VampPrior Mixture Model (VMM) as introduced in its original work [14]. This approach learns pseudo-inputs that act as cluster centroids in the latent space, with parameters optimized to represent the data distribution directly in the input space rather than only in the latent space. This provides a more flexible prior that encourages biologically meaningful clustering. The decoder maps the latent representation back to the data space by outputting the parameters of a Zero-inflated Negative Binomial (ZINB) distribution (mean, dispersion and zero-inflation probability). In this work the Negative Binomial distribution is also used to compare any meaningful differences in performance.

#### 4.1.2 First phase: Learning parameters of the prior distribution

The VAE training begins with the standard evidence lower bound (ELBO), augmented with additional penalization terms to structure the latent space according to biological groups. The loss function is defined as:

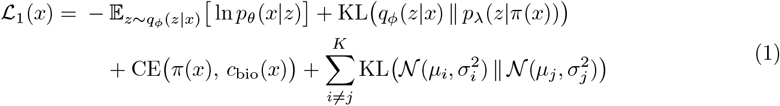

Here, *p*_*θ*_(*x*|*z*) is the decoder likelihood parameterized by *θ*, and *q*_*ϕ*_(*z*|*x*) is the encoder (approximate posterior) parameterized by *ϕ*. The variational prior distribution *p*_*λ*_(*z*|*π*(*x*)) is modeled as a Gaussian mixture with parameters 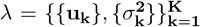, where **u**_**k**_ are pseudo-inputs mapped into the latent space and used as centroids of the *k*-th Gaussian component, and 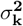 is its variance. The number of mixture components is denoted by **K**, that corresponds to the number of biological groups being modeled from the data.

The first term, 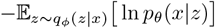, is the negative log-likelihood term of the ELBO and works as a reconstruction loss (i.e. how accurately the observed data *x* can be reconstructed from the latent representation *z*). The second term, KL (*q*_*ϕ*_(*z*|*x*)∥*p*_*λ*_(*z* |*π*(*x*))), is the KL-divergence term of the ELBO, which aligns the approximate posterior with the variational prior distribution. Because the prior is a mixture of Gaussians, the component selected for the KL calculation depends on the mixxing probablities *π*(*x*). The third term, CE (*π*(*x*), *c*_bio_(*x*)), is a **biological group assignment penalty**, implemented as the categorical cross-entropy between the inferred mixture probabilities *π*(*x*) and the one-hot encoded biological group *c*_bio_(*x*). This encourages samples in the same biological group to be mapped to the same latent component. Finally, the pairwise KL-divergence term, 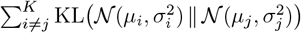, acts as a **clustering regularizer**, encouraging mixture components to be separated and therefore biological variability between groups to be preserved. Overall, this phase jointly optimizes reconstruction accuracy, prior structure, and biologically meaningful organization of the latent space.

#### 4.1.3 Second phase: Batch correction in the latent space

In the second phase, the prior parameters are set and unaltered for the rest of the training process: learned pseudo-inputs u_*k*_ and variance of each Gaussian component 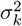 are fixed. A batch discriminator is then used to correct for any batch effect in the latent space. The discriminator takes latent points *z* as input and predicts their corresponding batch labels. The loss function for this phase is defined as:

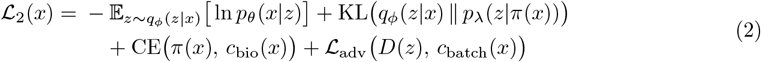

Where *D*(*z*) is the output of the adversarial network (discriminator), *c*_batch_(*x*) is the one-hot encoding of the batch label for *x*, and ℒ_adv_ *D*(*z*), *c*_batch_(*x*) is the adversarial loss that backpropagates through the encoder to encourage batch-invariant latent representations. The adversarial loss in this setup is defined as the negative cross-entropy between the discriminator output logits for *x* and *c*_batch_(*x*). In this setup, the discriminator learns to correctly predict batch labels, while the encoder is trained to maximize the discriminator’s error, effectively removing batch effect from the latent representation (similar to the objective used in the ABC framework [12]). Providing the batch label to the encoder and the decoder biases the model to reconstruct batch-specific features solely from the label; together with the adversarial objective, this encourages a batch-invariant latent *z* while preserving batch effects in the reconstructed output via the conditional decoder.

#### 4.1.4 Third phase: Decoder batch masking

The third phase starts with the encoder parameters being completely frozen, having only the decoder as a trainable component. The decoder’s loss function is limited to the negative log-likelihood:

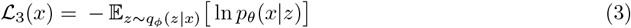

The idea here is to gradually mask the one-hot encoded batch labels previously provided to the decoder. Eventually, the decoder reconstructs the data solely from the batch-corrected clustered latent space. This procedure forces the decoder to eliminate any residual batch effects while preserving biological signals in the reconstructed data.

### 4.2 Simulated datasets generation

We generated two simulated datasets from a zero-inflated negative binomial (ZINB) distribution. Each scenario – (1) batch effect only, and (2) plus biological effect – consisted of 50 simulated replicates. Each replicate contained 200 samples and 1000 features, with samples assigned to two biological groups in equal proportion (50:50) and two batches in a 60:40 proportion. All draws from the distributions were performed with a fixed seed (42) to ensure reproducibility.

For each feature, the same ZINB parameters were used across all replicates. Specifically, for each feature, the dispersion was sampled as *r* ∼ Uniform(1, 3) and the baseline log-mean as *ℓ*_0_ ∼ *N* (2, 1); the baseline mean was then obtained by exponentiating *ℓ*_0_. Zero-inflation followed a two-tier scheme: 70% of features were designated as “highly sparse” with zero-probability *z* ∼ Uniform(0.8, 0.9), while the remaining 30% of features had *z* ∼ Uniform(0.1, 0.8).

#### 4.2.1 Batch effect only

For this scenario the baseline log-mean and variance were modified for samples from Batch 2, while no biological effect was introduced. While batch effects in practice can have broad impacts, we modeled a mixed scenario to evaluate both local and global artifacts. Specifically, for each observation in Batch 2, 10% of the features received an additive shift Δ_*batch*_ ∼ *N* (1, 2) to simulate feature-specific technical biases (a local batch effect). On top of this, batch-wide heteroskedastic adjustments were applied to all features: dispersion shifts δ_*disp*_ ∼ *N* (0, 0.2) and zero-inflation probability shifts δ_*zero*_∼ *N* (0, 0.2), representing global, variance-related technical differences.

For a sample in batch *b* ∈ {0, 1} (Batch 1 = 0, Batch 2 = 1), the adjusted mean is:

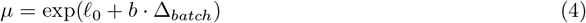

Next, the batch-dependent dispersion is given by:

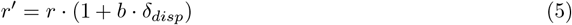

Finally, the batch-adjusted zero probability is given by:

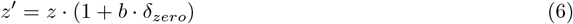

Where *z*’ ∈ [0, 1] to ensure a valid probability. In order to simulate correctly the ZINB distribution, the counts were sampled from NB(*µ, r*’) and a zero-mask was applied using Bernoulli(*z*’) to set the corresponding entries to zero.

#### 4.2.2 Batch and biological effect

For the simulated dataset with both batch and biological effects, the procedure extends the batch-only scenario. For 50% of the features in biological group Condition B, an additive log-scale effect was drawn as Δ_*bio*_∼ *N* (2, 2). The expected magnitude Δ_*bio*_ is larger than the batch shift, ensuring that the biological effect is the main source of additive variation. An interaction effect is also included to model biological-specific batch effects on 20% of features. This interaction shift was drawn from Δ_*int*_ ∼ *N* (3, 1) and applied additively on the log-scale mean only for samples belonging to both Batch 2 and Condition B.

For a sample in batch *b* ∈ {0, 1} (Batch 1 = 0, Batch 2 = 1) with condition *c* ∈ {0, 1} (Condition A = 0, Condition B = 1), the log-scale mean in:

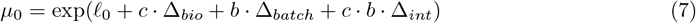

In addition, a per-sample indicator selects approximately 30% of features. For this subset (*u* ∈ {0, 1}), the mean is adjusted by a feature-wise factor:

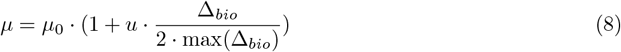

Dispersion and zero-inflation probabilities are obtained in the same way as in the batch-only simulations, with the counts drawn from the resulting ZINB distribution.

### 4.3 Case study datasets

#### 4.3.1 Anaerobic Digestion dataset

The anaerobic digestion dataset was used in a study benchmarking several batch effect assessment and correction approaches in microbiome data [5]. The data is publicly available through the GitHub repository linked to the work, where it was downloaded and converted to a comma-separated values (CSV) file for convenience. As mentioned before, it consists of 567 taxonomic groups and 75 samples, distributed across 5 batches (processing days) and 2 biological groups (treatment groups). The implementation is available as a demonstration in a Jupyter Notebook within the ABaCo repository.

#### 4.3.2 Inflammatory Bowel Disease dataset

The dataset is composed of two different BioProjects which were merged directly before pre-processing: one that studies the dynamics of microbiome functionality in IBD (accession: PRJNA389280) and another that includes multi-omics measurements from IBD patients in a longitudinal study (accession: PRJNA398089). Both studies are also available in MGnify (study IDs MGYS00002301 and MGYS00006120 respectively), from which the overall taxonomic assignments were downloaded. For consistency, only the genus-level taxa were retained, resulting in a dataset comprising 435 taxonomic groups. Metadata was retrieved from the Data Repository for Human Gut Microbiota (GMrepo) [26], where the phenotype identifiers were mapped to categorical labeles (i.e. D003425 →Crohn Disease, D003093 →Ulcerative Colitis, and D006262 →non IBD).

#### 4.3.3 DTU-GE sewage dataset

For the DTU-GE sewage dataset, we retrieved the taxonomic abundance tables directly from the MG-nify API for the analysis accessions listed in the study MGYS00001312. Data were obtained separately using the SSU rRNA taxonomic analysis from pipeline versions 4.1 and pipeline 3.0, capturing batch effect associated with computational framework differences. Counts were aggregated to the phylum level, and annotated with the geographic location of sampling to be used as a desirable co-founding variable (biological group). We retained samples from countries represented at least 15 times in the whole dataset, resulting in a total of 4 countries and 129 samples. Finally, taxa with zero counts across all samples were removed, yielding a final dataset of 162 taxonomic features.

### 4.4 Performance assessment

We used four metrics generally used to quantify batch-effect correction in sing-cell studies [27]: kBET [21], iLISI [22], batch ASW [23] and batch ARI [24] - to assess batch effect removal, where values closer to 1 indicate better performance.

We applied PCoA using the Aitchison distance to compare clustering by batch and biological group both before and after correction. To quantify the variance explained by batch and biological factors, we performed a Permutational Multivariate Analysis of Variance (PERMANOVA) test with 999 permutations, reporting the R^2^ attributable to each factor independently [28].

For the robustness analysis, we track the 5 most abundant taxa in each case study dataset after correction using ABaCo by reporting their mean relative abundance across all runs. We report the mean and variance for each biological group to verify the consistency of ABaCo’s inference in each iteration. Additionally, we applied the Kruskal-Wallis test [28] to all taxonomic features to identify significant differences among biological groups before and after correction. For each taxon, we computed the K-W statistic and its p-value, adjusting for multiple testing across taxa using the Benjamini-Hochberg false discovery rate. Taxa with an adjusted p-value below 0.05 were considered significantly differentially abundant in at least one biological group (see Supplementary Figure 12).

State-of-the-art methods for batch effect correction use either normalized counts (Batch Mean Centering (BMC) [5], ComBat [6], limma [7], PLSDA-batch [10]) or raw counts (ComBat-seq [29], ConQuR [9] and ABaCo (ours)). Because ABaCo implements a generative model, all analyses were repeated for 50 training runs, and we report both mean performance and variability to quantify the consistency of the model.

### 4.5 Experimental setup

The model architecture and training hyperparameters were kept consistent across all datasets. the encoder consists of 512, 256 and 128 neurons, respectively, and the decoder mirrors this with three layers of 128, 256 and 512 neurons. The latent space dimension was set to 16 for all datasets except the IBD case study, where 32 dimensions were used. ReLU activations were applied between each layer. The batch discriminator network consists of three fully connected layers (256, 128 and 64 neurons) with ReLU activations. A summary of the training hyperparameters used in each dataset is provided in the Supplementary Table 1. The epochs for each phase (1st, 2nd, 3rd) and the values of all learning rates used—Phases lr.: VAE learning rates for each phase; Disc. lr.: learning rate used when training the discriminator during the second phase; Adv. lr.: learning rate used to backpropagate the adversarial loss to the encoder—are define in here. The weights for each term on the losses are also mentioned at the Supplementary Table 2. NLL: negative log-likelihood term of the ELBO; KL-div: KL-divergence between prior *p*_*λ*_(*z*) and posterior *q*_*ϕ*_(*z* | *x*) distributions; Bio. penalty: cross-entropy used for the biological group assignment penalty; Clust. penalty: pairwise KL-divergence of prior Gaussian components used for the cluster overlap penalty.

Model training and evaluation were performed on a dual-socket Intel Xeon Gold 6226R server running in x86 64 mode with VT-x virtualization enabled. Each socket contains 16 physical cores with Hyper-Threading (2 threads per core), yielding a total of 64 logical CPUs. The system is equipped with 1 TiB of RAM and runs Debian 12 “Bookworm” with Linux kernel 6.1.0-37-amd64. For GPU acceleration, two NVIDIA Tesla V100S-PCIE cards (32 GiB each) were installed, using NVIDIA driver 565.57.01 and CUDA 12.7. The compute performance is summarized in Supplementary Table 3.

### 4.6 ABaCo Python Library

All the code for ABaCo is openly available on GitHub (https://github.com/Multiomics-Analytics-Group/abaco) under the MIT license, and is distributed as a Python package in PyPI (https://pypi.org/project/abaco/). The documentation is provided at ReadtheDocs (https://mona-abaco.readthedocs.io/) where there are several tutorials exemplifying how to use ABaCo.

## Supplementary material

**Supplementary Table 1:** Training schedule for simulated and case study datasets. The table lists the number of epochs for each training phase (Phase 1–3), the VAE learning rate in each phase (Phase lr.), the batch-discriminator learning rate (Disc. lr.), and the encoder adversarial learning rate used during discriminator updates (Adv. lr.).

| **Dataset** | **Epochs**<br>(1st, 2nd, 3rd) | **Phases lr.**<br>(1st, 2nd, 3rd) | **Disc. lr.** | **Adv. lr.** |
| --- | --- | --- | --- | --- |
| Anaerobic digestion | 2000 | 1e-3 | 1e-3 | 1e-3 |
|  | 2000 | 1e-3 |  |  |
|  | 1000 | 1e-6 |  |  |
| Inflammatory Bowel Disease | 4000 | 1e-4 | 1e-6 | 1e-6 |
|  | 1000 | 1e-6 |  |  |
|  | 2000 | 1e-6 |  |  |
| DTU-GE sewage | 4000 | 2e-4 | 1e-6 | 1e-6 |
|  | 1000 | 1e-6 |  |  |
|  | 1000 | 1e-7 |  |  |
| Simulated data<br>Batch effect only | 2000 | 1e-4 | 1e-5 | 1e-5 |
|  | 2000 | 1e-5 |  |  |
|  | 1000 | 1e-7 |  |  |
| Simulated data<br>Batch and biological<br>effect | 5000 | 2e-4 | 1e-7 | 1e-7 |
|  | 2000 | 1e-6 |  |  |
|  | 1000 | 1e-6 |  |  |

**Supplementary Table 2:**
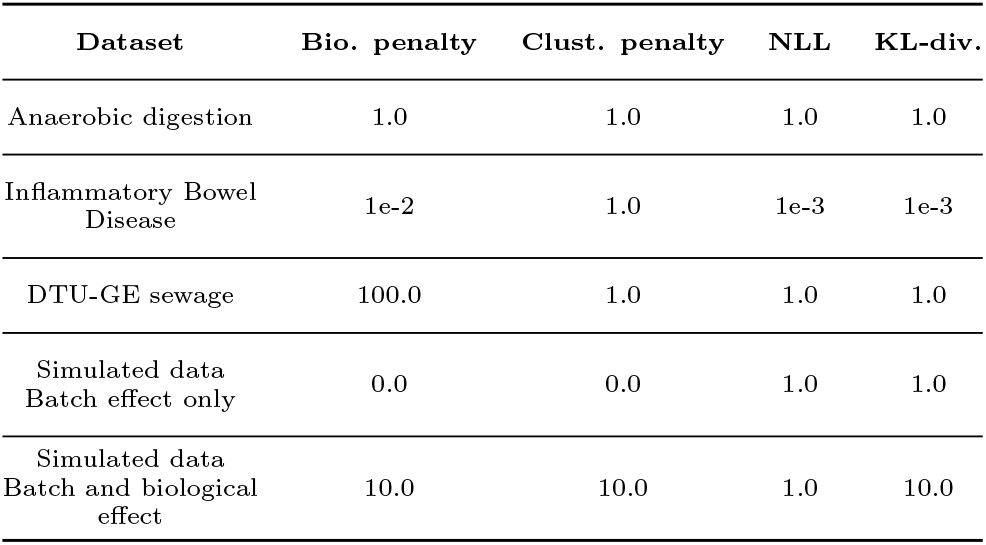
Weights used for the loss function in simulated and case study datasets. The table shows the weight of the biological preservation penalty (Bio. penalty), the cluster overlap penalty (Clust. penalty), the negative log-likelihood (NLL), and the KL-divergence (KL-div.) terms of the ELBO.

**Supplementary Table 3:** Compute performance of ABaCo with different decoder output distribution **ZINB**: Zero-inflated Negative Binomial; **NB**: Negative Binomial.

| Dataset | Samples | Taxa | Decoder | Compute Metrics |  |  |
| --- | --- | --- | --- | --- | --- | --- |
|  |  |  |  | CPU Time (s) | Single GPU Util. (%) | Peak GPU Mem. (GiB) |
| Anaerobic Digestion | 75 | 567 | ZINB | 77.2 | 26 | 0.07 |
|  |  |  | NB | 62.8 | 24 | 0.06 |
| Inflammatory Bowel Disease | 524 | 435 | ZINB | 117.1 | 19 | 0.05 |
|  |  |  | NB | 115.3 | 17 | 0.05 |
| DTU-GE sewage | 129 | 162 | ZINB | 112.9 | 24 | 0.04 |
|  |  |  | NB | 103.7 | 22 | 0.04 |
| Simulated data:<br>Batch effect only | 200 | 1000 | ZINB | 75.7 | 27 | 0.10 |
|  |  |  | NB | 71.9 | 24 | 0.08 |
| Simulated data:<br>Batch and biological effect | 200 | 1000 | ZINB | 121.1 | 27 | 0.10 |
|  |  |  | NB | 109.2 | 25 | 0.08 |

**Supplementary Figure 1:**
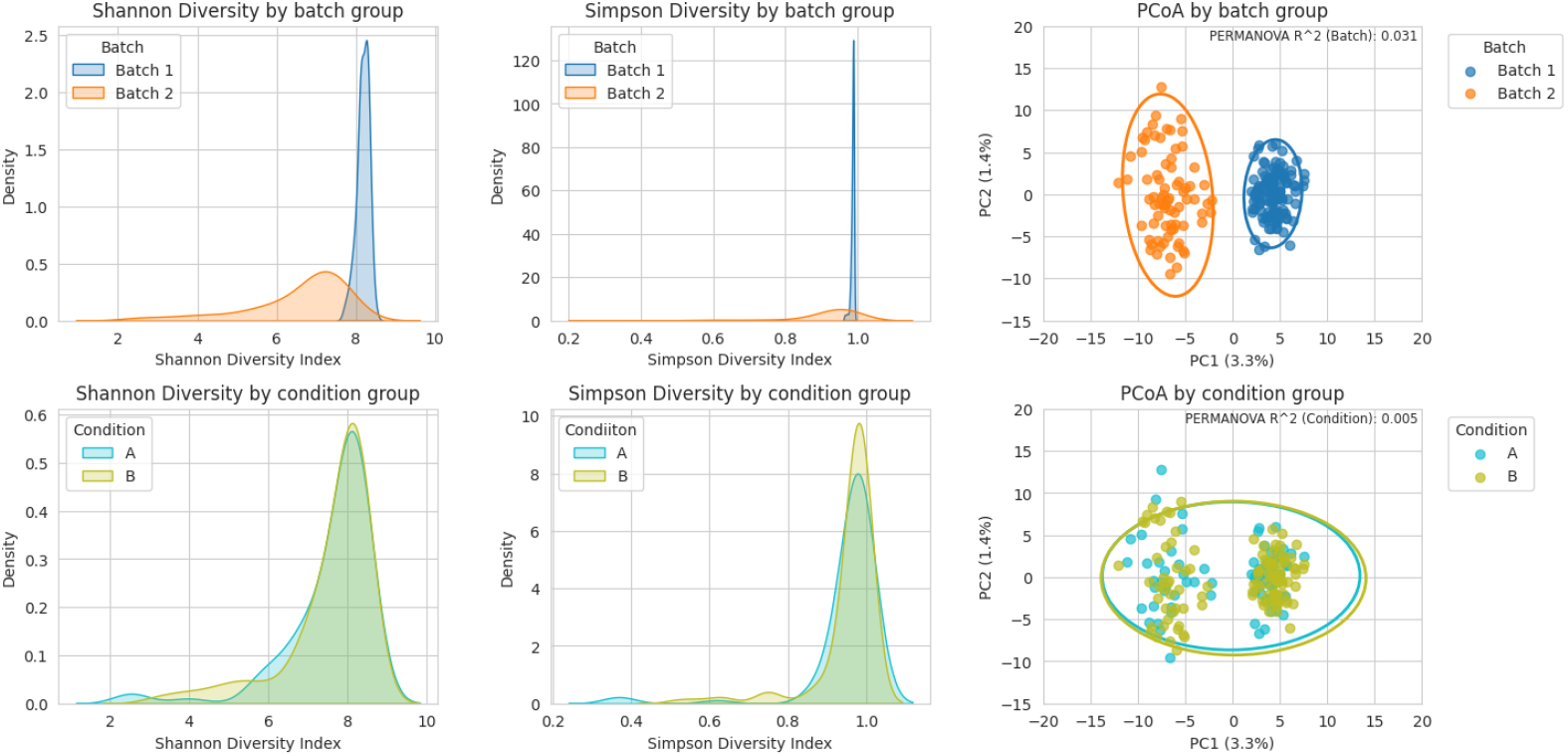
Kernel density estimates of Shannon and Simpson alpha diversity (left, center) and PCoA on Aitchison distances (right), shown by batch (top) and biological group (bottom) for the simulated dataset with batch effect only.

**Supplementary Figure 2:**
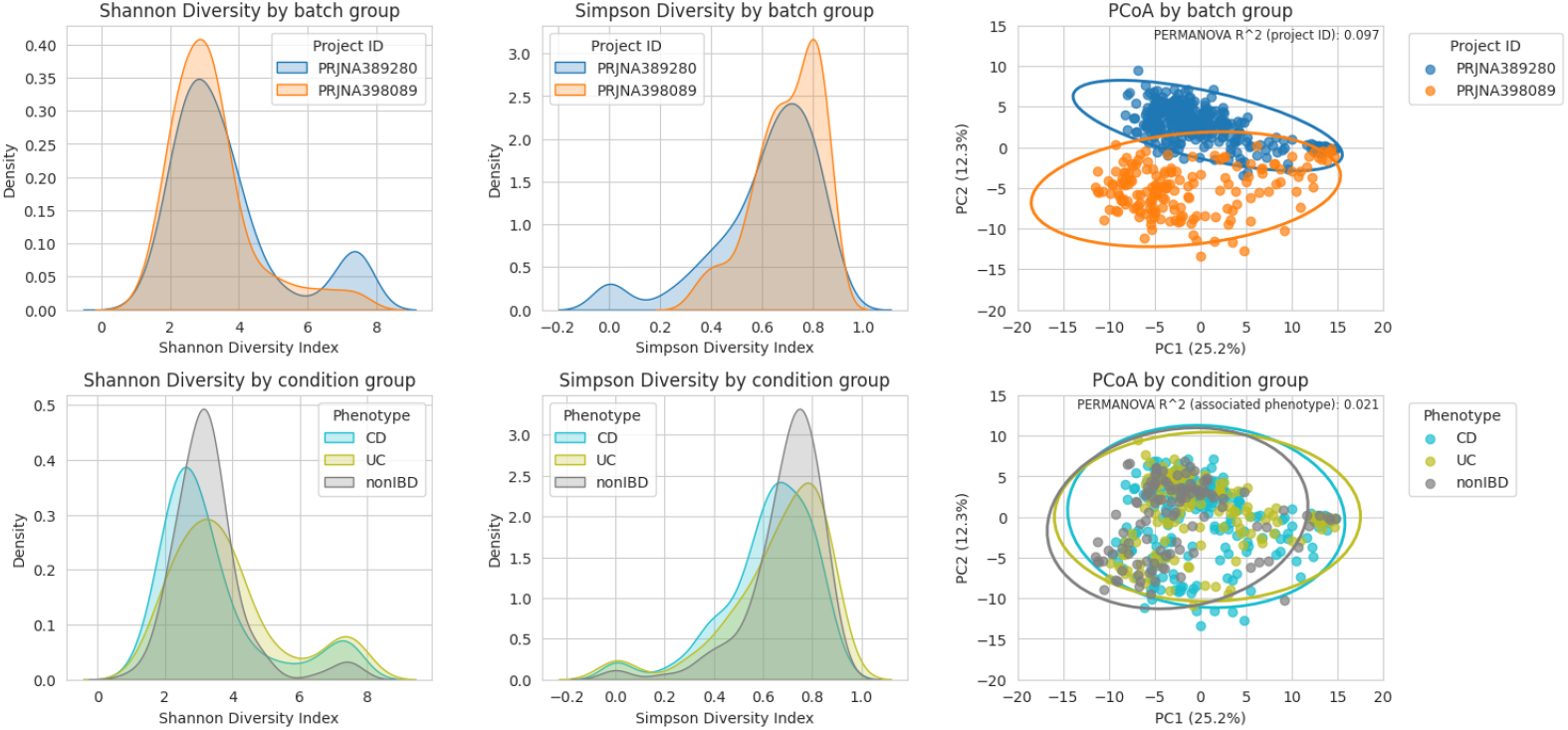
Kernel density estimates of Shannon and Simpson alpha diversity (left, center) and PCoA on Aitchison distances (right), shown by batch (top) and biological group (bottom) for the IBD case study dataset.

**Supplementary Figure 3:**
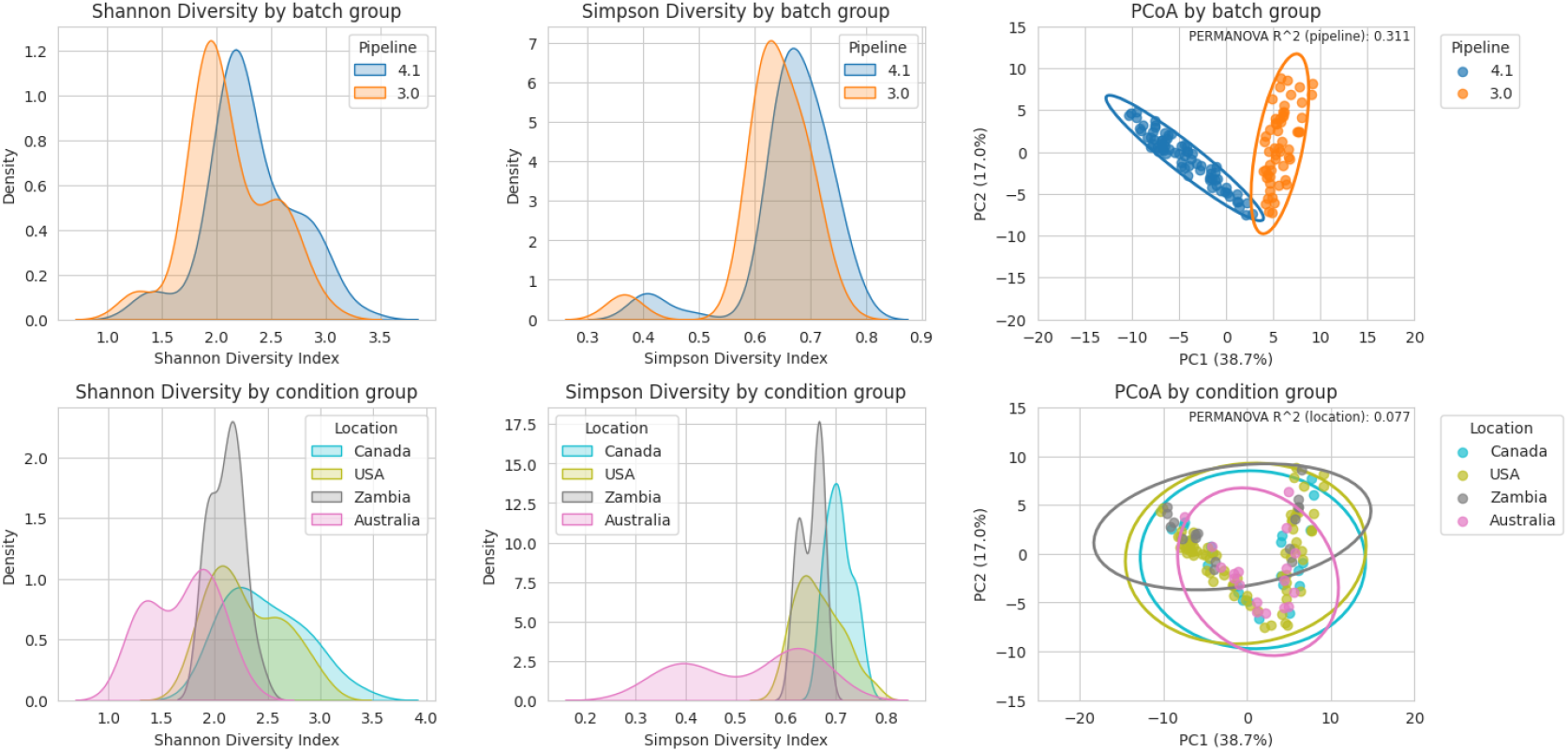
Kernel density estimates of Shannon and Simpson alpha diversity (left, center) and PCoA on Aitchison distances (right), shown by batch (top) and biological group (bottom) for the DTU-GE sewage case study dataset.

**Supplementary Figure 4:**
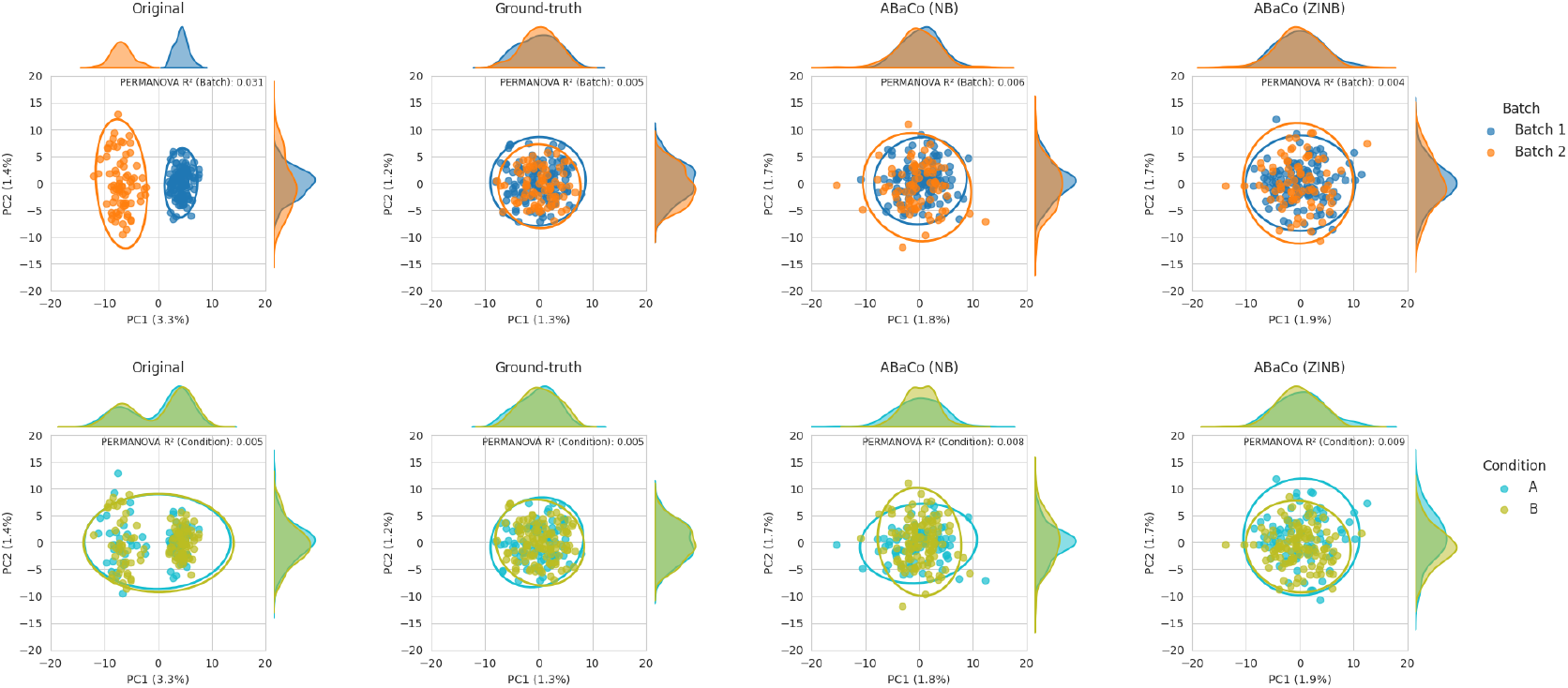
Principal Coordinates Analysis (PCoA) using Aitchison distance on the simulated data containing batch effect only corrected with ABaCo; the ground-truth PCoA is shown for reference.

**Supplementary Figure 5:**
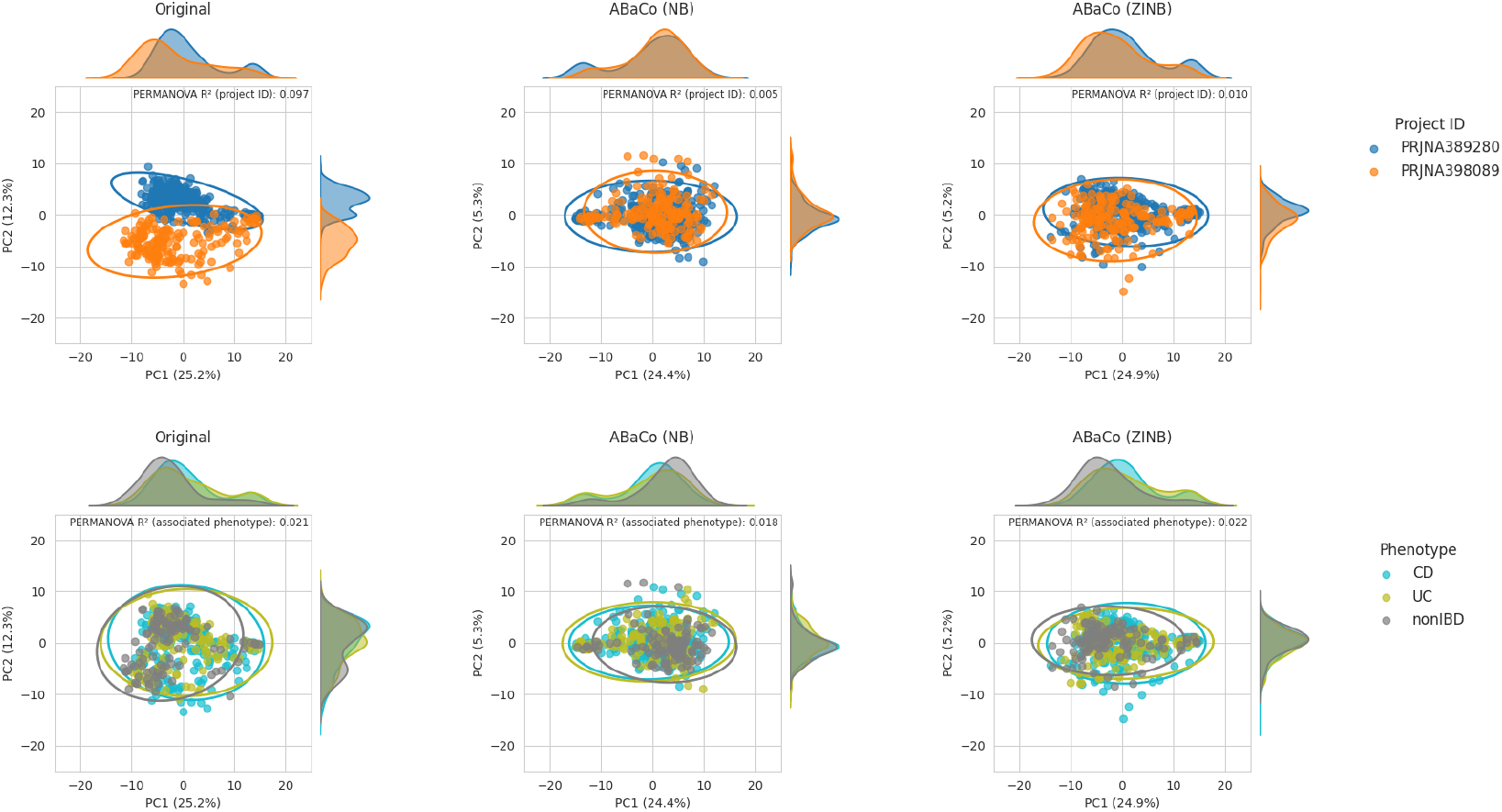
Principal Coordinates Analysis (PCoA) using Aitchison distance on the IBD case study dataset corrected with ABaCo.

**Supplementary Figure 6:**
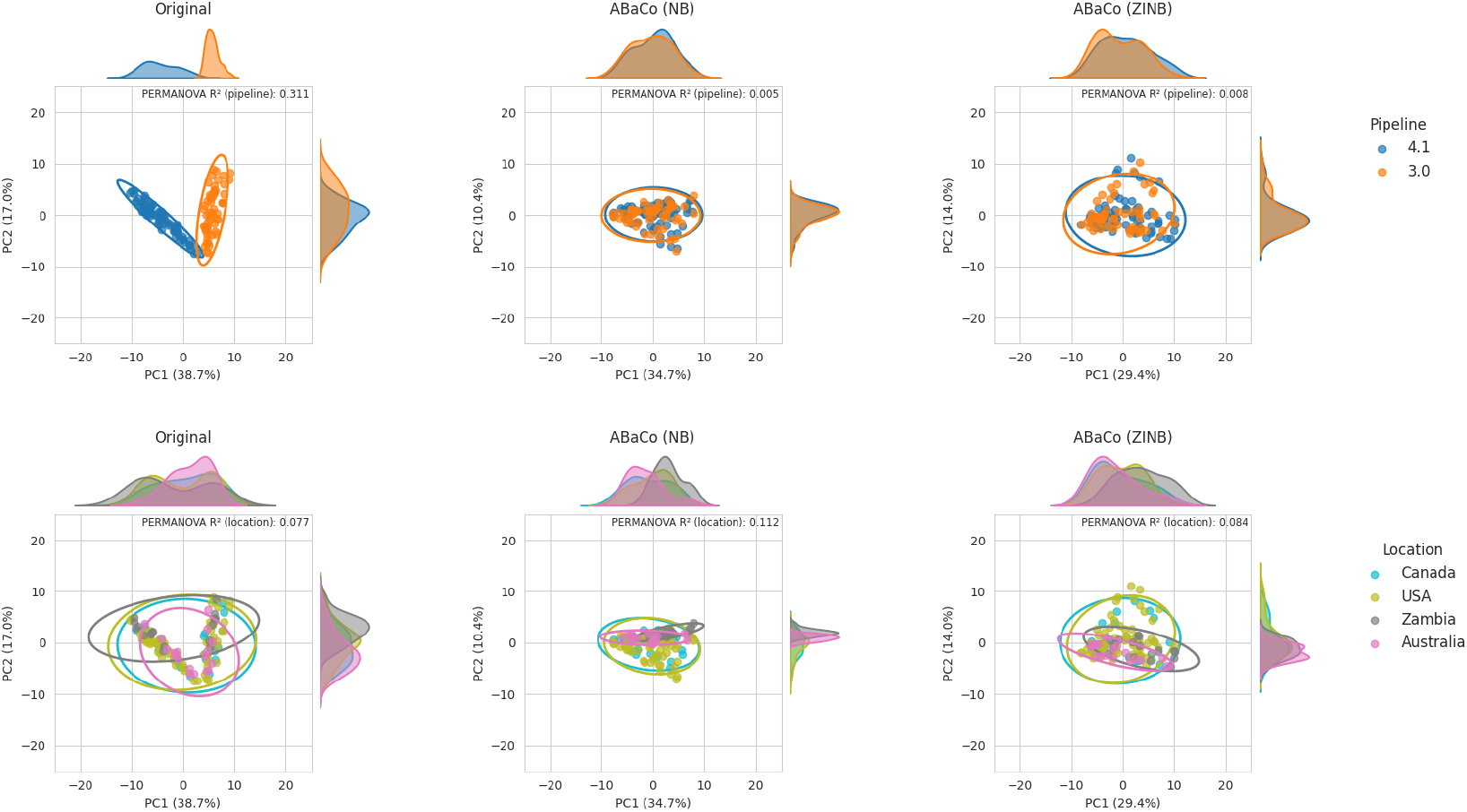
Principal Coordinates Analysis (PCoA) using Aitchison distance on the DTU-GE sewage case study dataset corrected with ABaCo.

**Supplementary Figure 7:**
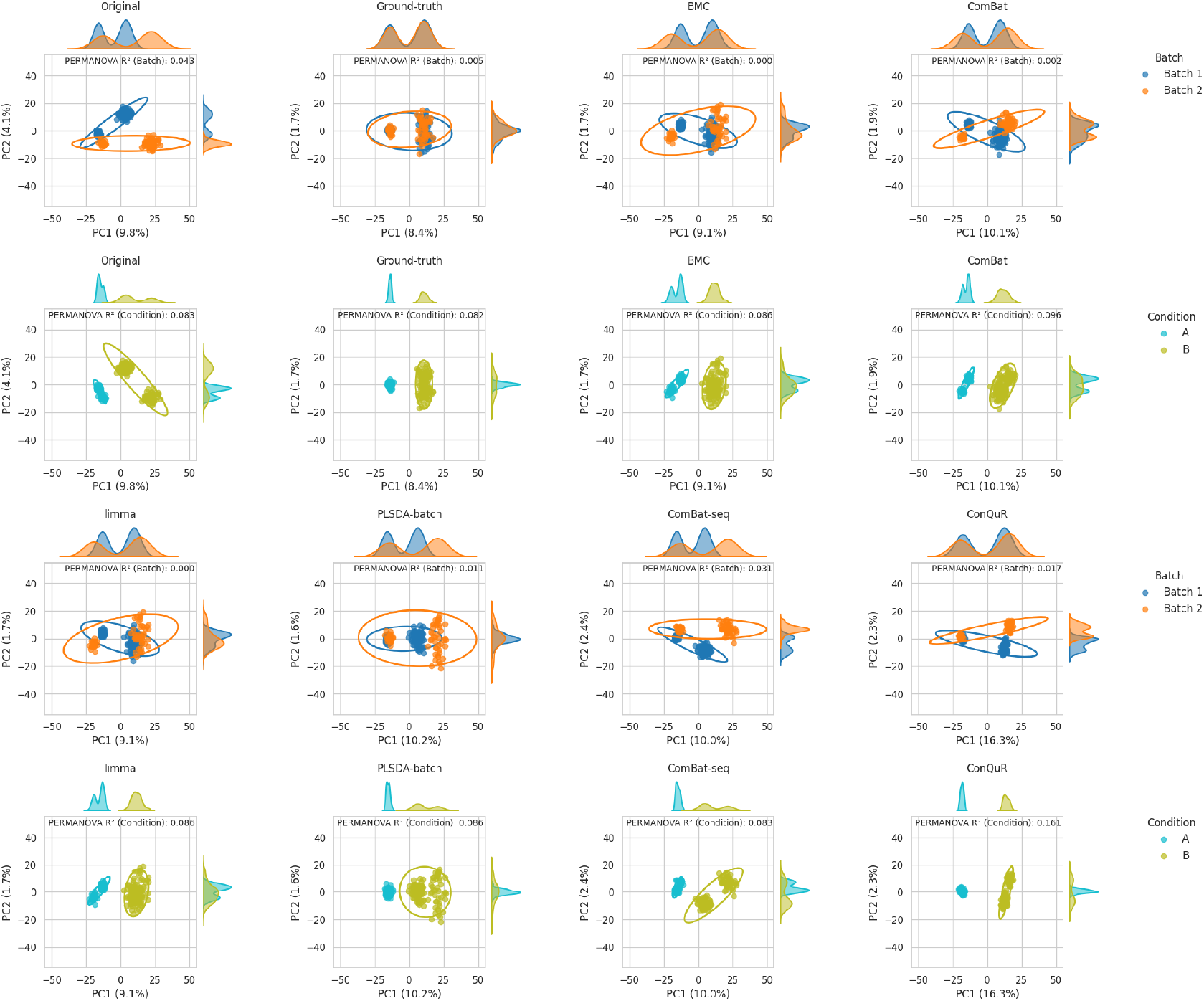
Principal Coordinates Analysis (PCoA) using Aitchison distance on the simulated data containing both batch and biological effect corrected with state-of-the-art methods; the ground-truth PCoA is shown for reference.

**Supplementary Figure 8:**
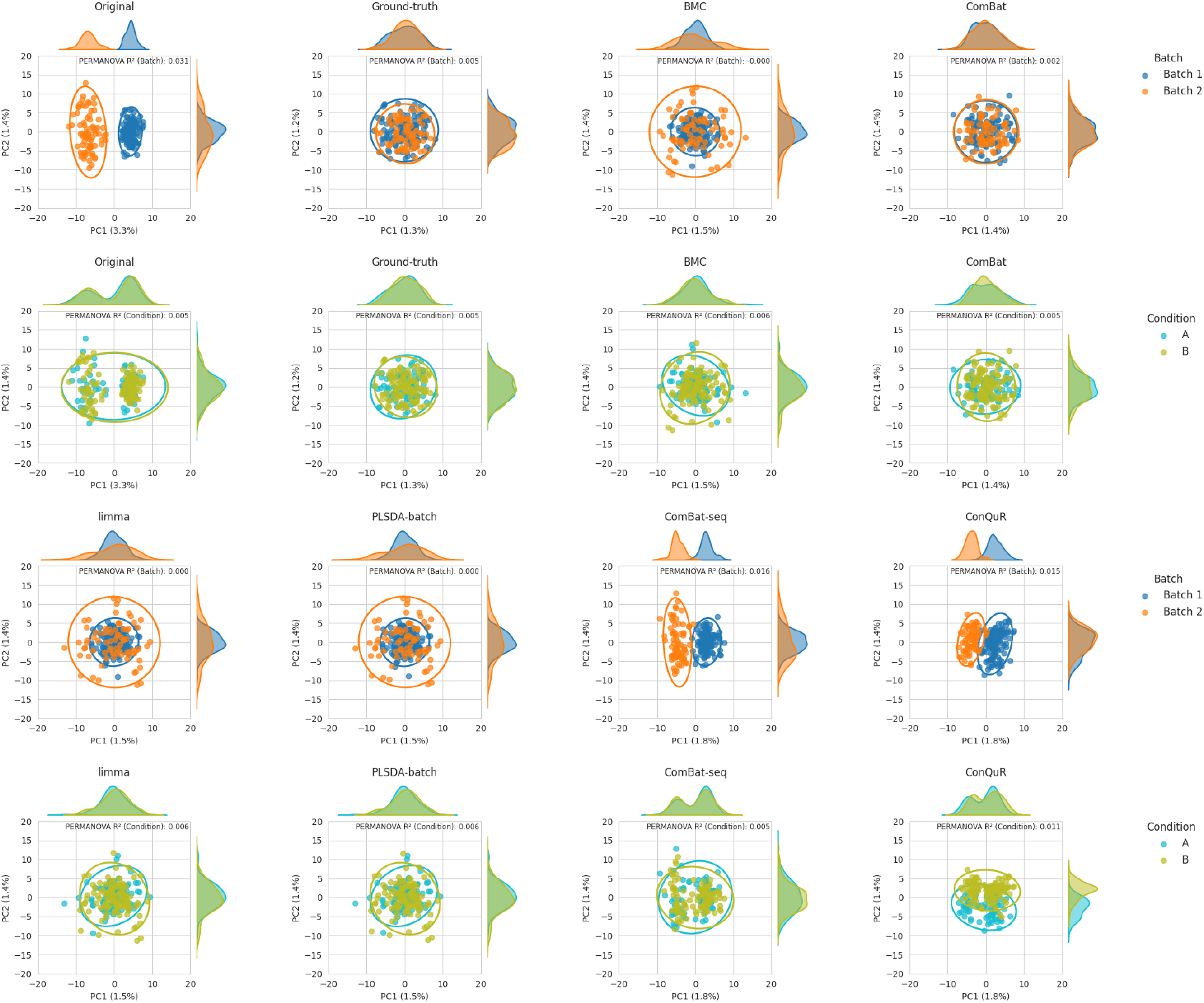
Principal Coordinates Analysis (PCoA) using Aitchison distance on the simulated data containing batch effect only corrected with state-of-the-art methods; the ground-truth PCoA is shown for reference.

**Supplementary Figure 9:**
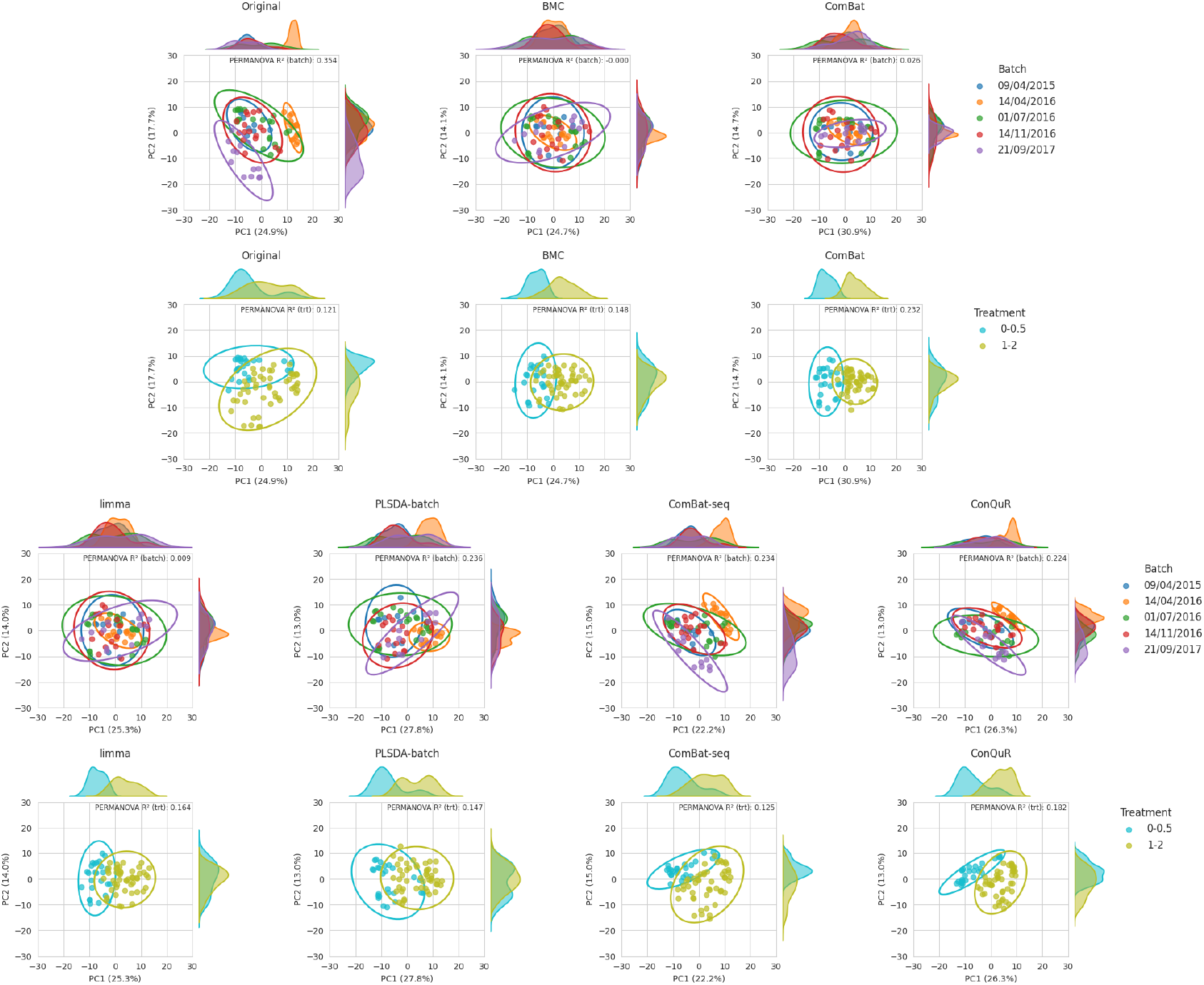
Principal Coordinates Analysis (PCoA) using Aitchison distance on the anaerobic digestion case study dataset corrected with state-of-the-art methods.

**Supplementary Figure 10:**
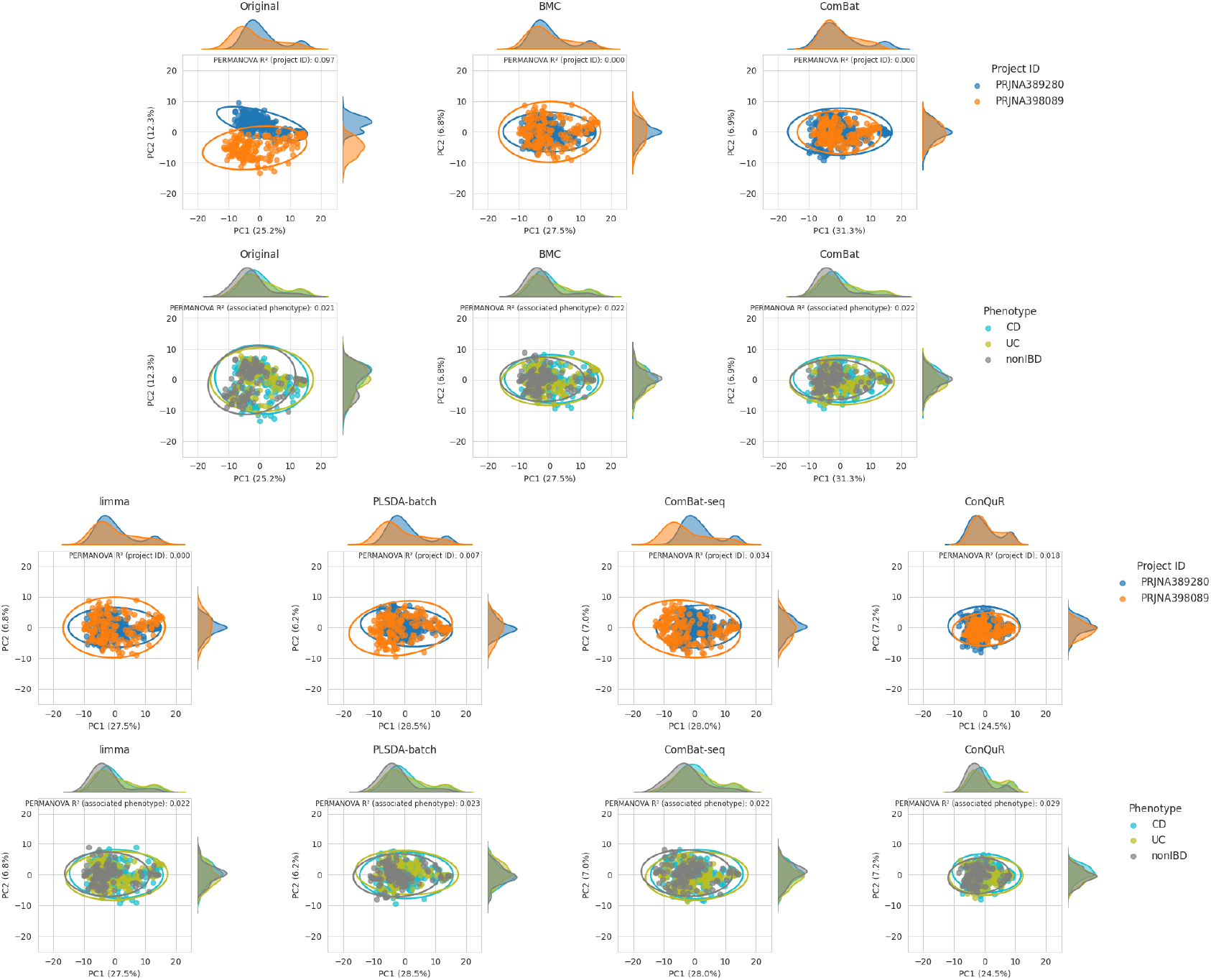
Principal Coordinates Analysis (PCoA) using Aitchison distance on the IBD case study dataset corrected with state-of-the-art methods.

**Supplementary Figure 11:**
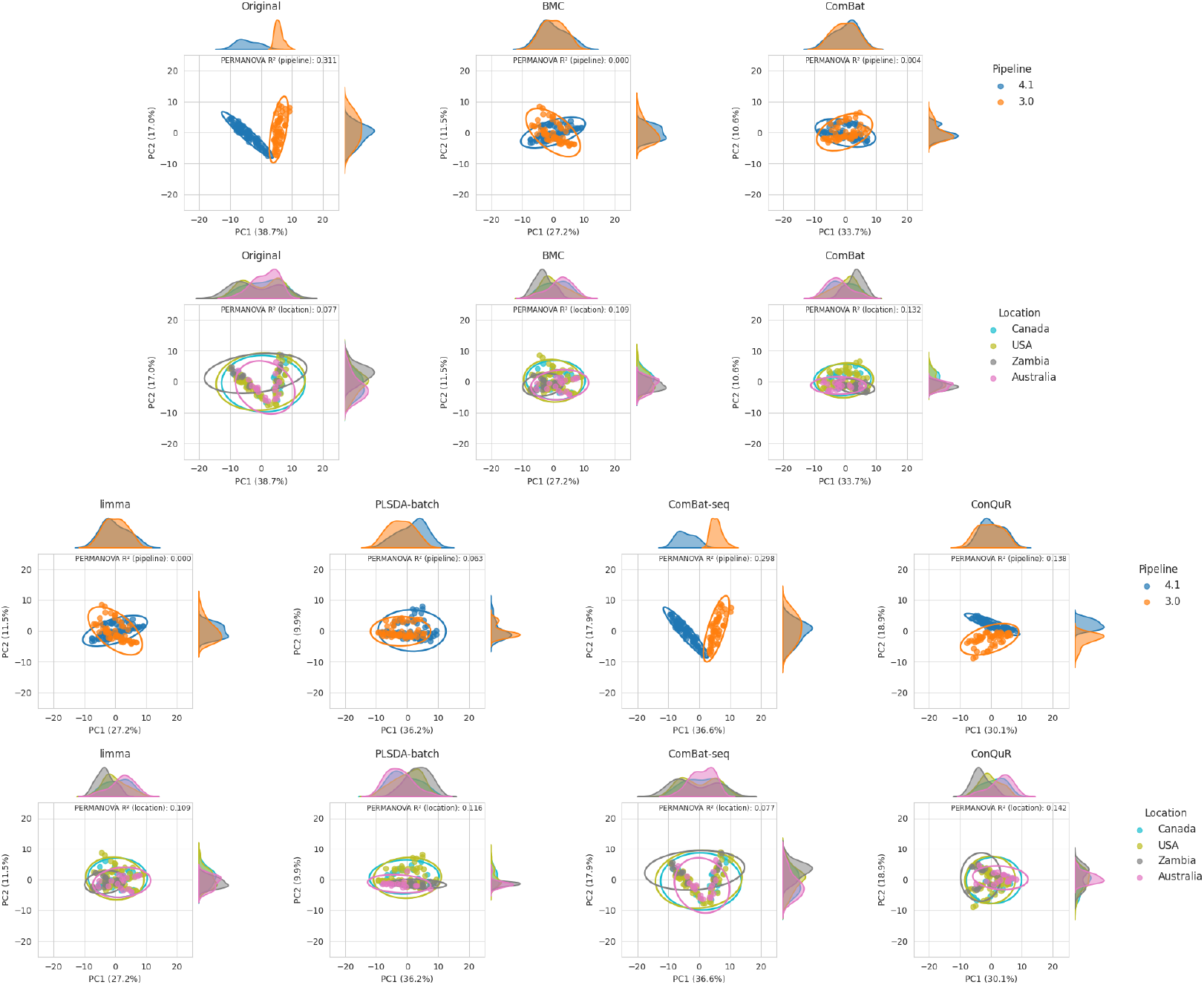
Principal Coordinates Analysis (PCoA) using Aitchison distance on the DTU-GE sewage case study dataset corrected with state-of-the-art methods.

**Supplementary Figure 12:**
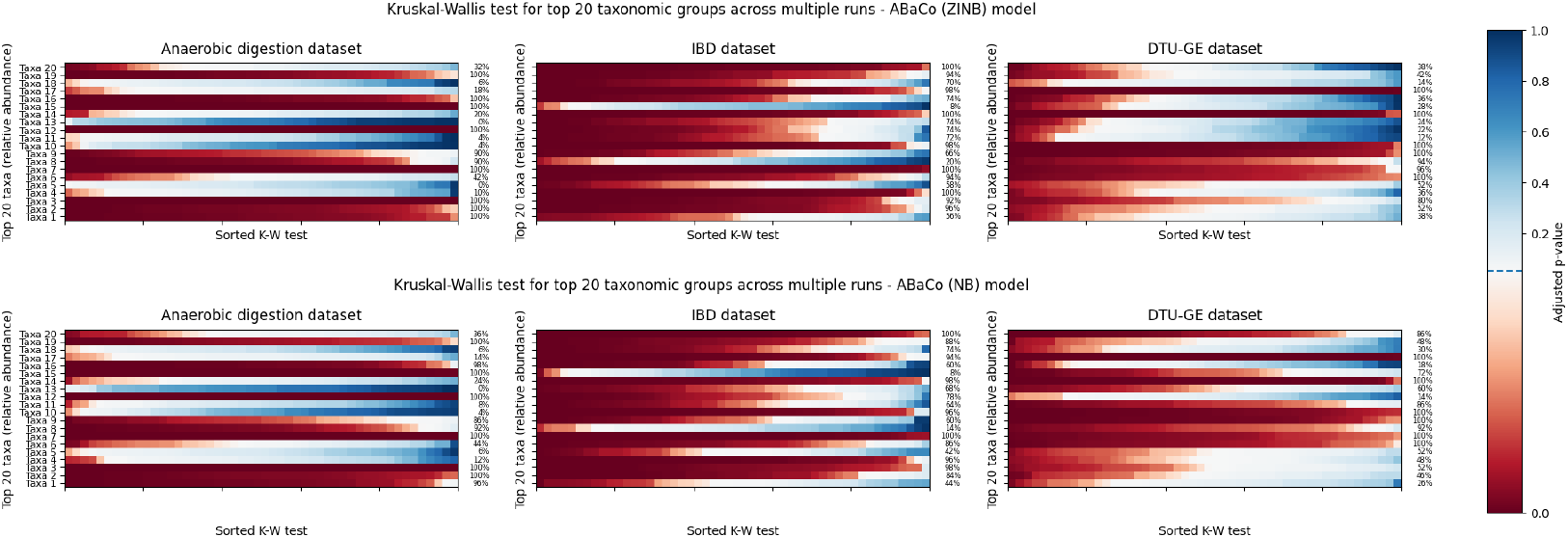
Kruskal-Wallis test results in most abundant taxa for each case study dataset corrected with ABaCo.

**Supplementary Figure 13:**
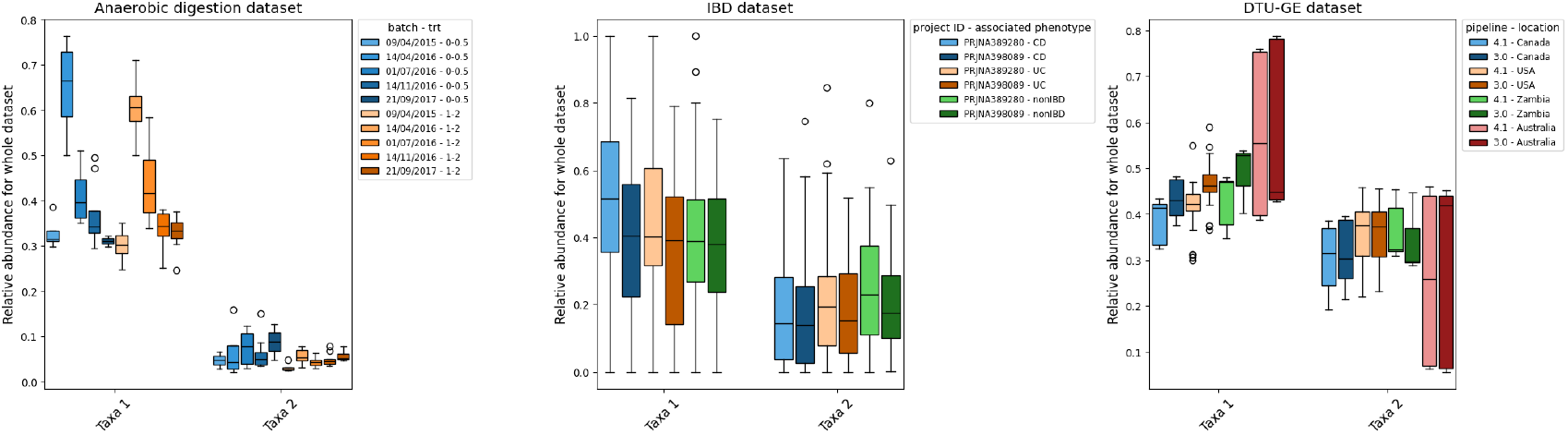
Relative abundance of the two main taxa for all case study datasets stratified by biological and batch groups.

